# Heteromeric RNP assembly at LINEs controls lineage-specific RNA processing

**DOI:** 10.1101/297853

**Authors:** Jan Attig, Federico Agostini, Clare Gooding, Aarti Singh, Anob M Chakrabarti, Nejc Haberman, Warren Emmett, Christopher WJ Smith, Nicholas M Luscombe, Jernej Ule

**Affiliations:** The Francis Crick Institute, Midland Road 1, Kings Cross, London NW1 1AT; Department of Molecular Neuroscience, UCL Institute of Neurology, Queen Square, London, WC1N 3BG, UK; Department of Biochemistry, University of Cambridge, Tennis Court Road, Cambridge, CB2 1QW, UK; Department of Comparative Biomedical Sciences, The Royal Veterinary College, Royal College Street, London NW1 0TU, UK; Department of Genetics, Environment and Evolution, UCL Genetics Institute, Gower Street, London WC1E 6BT, UK; Okinawa Institute of Science & Technology Graduate University, 1919-1 Tancha, Onna-son, Kunigami-gun, Okinawa 904-0495, Japan

**Keywords:** splicing, pre-mRNA processing, LINE repeats, MATR3, PTBP1

## Abstract

It is challenging for RNA processing machineries to select exons within long intronic regions. We find that intronic LINE repeat sequences (LINEs) contribute to this selection by recruiting dozens of RNA-binding proteins (RBPs). This includes MATR3, which promotes binding of PTBP1 to multivalent binding sites in LINEs. Both RBPs repress splicing and 3’ end processing within and around LINEs, as demonstrated in cultured human cells and mouse brain. Notably, repressive RBPs preferentially bind to evolutionarily young LINEs, which are confined to deep intronic regions. These RBPs insulate both LINEs and surrounding regions from RNA processing. Upon evolutionary divergence, gradual loss of insulation diversifies the roles of LINEs. Older LINEs are located closer to exons, are a common source of tissue-specific exons, and increasingly bind to RBPs that enhance RNA processing. Thus, LINEs are hubs for assembly of repressive RBPs, and contribute to evolution of new, lineage-specific transcripts in mammals.

## INTRODUCTION

Human introns are replete with sequences that resemble splice sites and polyA sites, creating a demand for mechanisms that can help processing machineries distinguish true from cryptic RNA processing sites (Semlow and Staley, 2012, Sibley et al., 2016). Inappropriate recognition of such sites initiates inclusion of cryptic exons, which can disrupt gene expressing by changing the reading frame, introducing premature stop codons, and decreasing transcript stability. Mutations that activate splicing of cryptic exons therefore cause a number of hereditary human diseases (Vorechovsky, 2010, Sibley et al., 2016). It is well understood that exon definition mechanisms maintain splicing fidelity by combinatorial recognition of 3’ and 5’ splice sites and exonic enhancer sequences. Moreover, RNA-binding proteins (RBPs) can contribute to splicing fidelity by directly repressing the cryptic splice sites of RNA processing (Reed, 2000). Therefore, mutations disrupting their binding sites can activate cryptic exons and cause disease (Eom et al., 2013, Vorechovsky, 2010). However, the RBPs that promote splicing fidelity by assembling over deep intronic regions, and the elements that coordinate assembly of ribonucleoprotein complexes (RNPs) across introns are yet to be fully identified.

The human genome contains over 1.5 million LINE repeats, retrotransposons, many of which are located in introns. The two most common LINE repeat families in mammals are L1 and L2, which first inserted before the split of monotremata and therians approximately 200 million years ago (O’Leary et al., 2013), and new subfamilies have amplified in bursts ever since (Huang et al., 2010). The consensus L1 sequence contains strong cryptic splice sites in both sense and antisense orientation (Belancio et al., 2006, Merkin et al., 2015). The effect of intragenic LINEs on splicing has been studied mainly in the context of pathological conditions, in which intronic LINE insertions disrupt expression of their host gene. For instance, creation of new cryptic LINE-derived exons disrupt expression of the *CYBB* gene in individuals with chronic granulomatous disease (Meischl et al., 2000) and the *DMD* gene in X-linked dilated cardiomyopathy (Yoshida et al., 1998), and a less well characterised intronic LINE insertion disrupts expression of *XRP2* in X-linked retinitis pigmentosa 2 (Schwahn et al., 1998). Yet, only 60-80 LINEs were found capable of retrotransposition in the human genome, and account for all the *de novo* LINE insertions observed in human populations or *in vitro* (Beck et al., 2010, Brouha et al., 2003). In the remaining LINEs, mutations have disrupted the ability to retrotranspose, and most are degenerated and truncated compared to the consensus sequence. In many genes, such degenerated LINEs form part of their introns. Several RBPs are known to bind active LINEs and thereby interfere with their retrotransposition (Goodier et al., 2013, Taylor et al., 2013, Goodier et al., 2012), but the RBPs binding intronic LINEs, and the regulatory potential of these LINEs, are poorly understood.

Here, we surveyed iCLIP and eCLIP data to identify 28 RBPs that have enriched binding to intragenic LINEs. Notably many of these RBPs, including MATR3 and PTBP1, primarily bind to deep intronic regions i.e. more than >500nt away from any known exon. MATR3 promotes binding of PTBP1 to LINEs at clusters rich in CU-binding motifs and, together, the two RBPs create a repressive environment by blocking the recognition of cryptic polyA-sites and splice sites. Analysis of distinct evolutionary classes of intragenic LINEs demonstrated that repressive RBPs are most enriched on younger, primate-specific L1 elements, which are depleted in the vicinity of exons. In contrast, the binding of repressive RBPs is decreased on evolutionary older LINEs, especially those preserved across mammals, with a concomitant increase in binding of RBPs that are involved in recognition of 3’ and 5’ splice sites, and polyA sites. These older LINEs are located in closer proximity to exons, and are a source of new mammalian exons, with higher inclusion levels and differential regulation across human tissues. Thus, while most LINEs recruit repressive RBPs to insulate the deep intronic regions from RNA processing, many older LINEs have started to escape from this repression. Hence, LINEs facilitate evolutionary innovations in the emergence of mammal-specific transcripts.

## RESULTS

### LINE-derived sequences recruit dozens of RBPs to deep intronic regions

The primary goal of our study was to identify repressive RBPs that assemble over deep intronic regions in a coordinated manner to distinguish genuine exons from other exon-like sequences that are present within introns. Because many cryptic exons arise from retrotransposable elements, which are pervasive in introns (Smit et al., 1996-2010a, Deininger and Batzer, 2002), we hypothesised that retrotransposons might be RNP assembly points in deep intronic regions. We focused on LINEs, because they constitute the largest proportion of intronic sequence, and are generally excluded from mRNAs but not pre-mRNA, as is evident from their presence in nuclear but not cytoplasmic transcriptomes in HeLa, K562 and HepG2 cell lines, and from their depletion in nuclear polyA+ compared to polyA-RNA (Figure 1A). To identify RBPs that bind to L1-derived sequences, we examined iCLIP data for 17 RBPs published by our laboratory, and eCLIP data from K562 and HepG2 cells for 112 RBPs available from ENCODE (Sloan et al., 2016, Van Nostrand et al., 2017). We ranked these RBPs by the proportion of crosslink events mapping to sense or antisense L1 elements (Figure 1B), as well as by their propensity to bind to deep intronic regions (Figure 1C). Overall, find that RBPs with most enrichment on L1 elements also rank highly for deep intronic binding.

**Figure 1:**
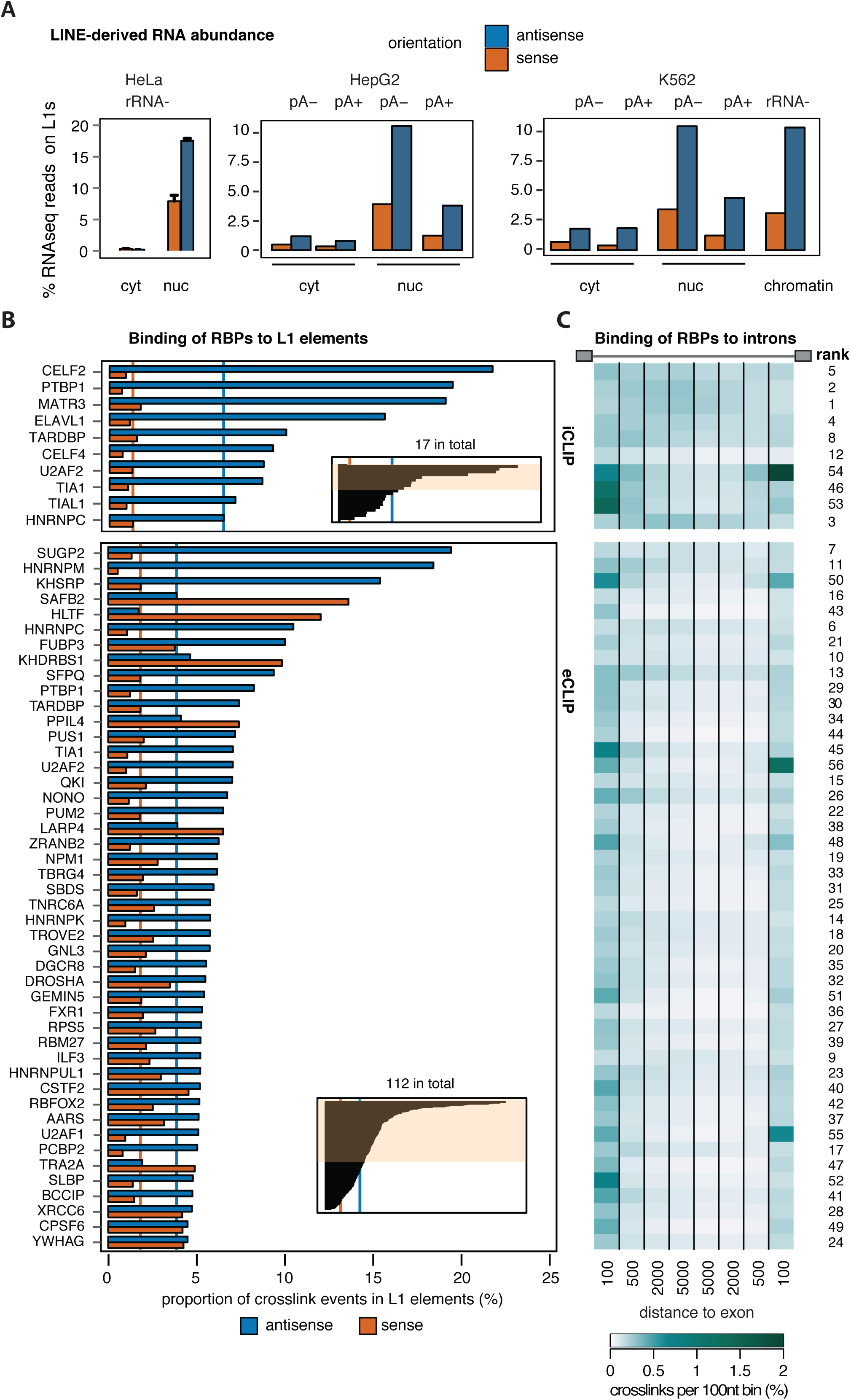
LINEs are binding platforms for a set of RBPs. (A) Estimate of abundance of L1-sequences in cytoplasmic and nuclear RNA fractions from HeLa, K562 and HepG2 cells. Strand-specific RNAseq was used to quantify abundance of L1 in sense and antisense (colored in orange and blue), relative to number of mapped reads. Data is split for libraries made from polyA-, polyA+ or rRNA-RNA. Data for K562 and HepG2 is from the ENCODE consortium. Data for HeLa is from replicates, and bargraph show mean ± s.d.m. (B) Frequency of L1 repeat sequences among the bound RNA sequences of a panel of RBPs. For each RBP, all cDNAs recovered in an iCLIP or eCLIP experiment were counted if they mapped at least partially to a L1 element. Since e/iCLIP is strand-specific, binding to LINEs transcribed in sense or in antisense was quantified separately, coloured in orange and blue. The orange and blue lines indicate the average binding across all RBPs (median). The iCLIP data was derived either from HeLa cells or from HEK293 FlpIN cells, and the eCLIP data from K562 and HepG2 cells. This information and the full data set is available in Suppl. Table 1, together with the source of each data set. For visualisation, replicates were averaged and only data from one cell line is shown. (C) Binding to introns of at least 7kb size was analysed in 100nt bins up to 5kb upstream and downstream of the exon, and quantified in percent relative to the total number of mapped reads. Data is shown for the first 100nt and as an average of bins 101-500nt, 501-2000nt and 2001-5000nt. A rank for deep intronic binding is given based on the average of the first 100nt of either splice site and average binding in the 20001-5000nt window.

While many RBPs show binding in deep intronic regions, MATR3 and PTBP1 ranked highest in iCLIP data (Figure 1C). Both RBPs have strong enrichment on L1s: 19% of MATR3 and PTBP1 iCLIP crosslink events were in antisense L1 repeats, which is a strong overrepresentation compared to the median of 6.4% across all examined iCLIP data (Figure 1B and SupplTable1). This agrees with a previous study that also found enrichment of PTBP1 iCLIP reads in LINEs compared to a genomic null model (Kelley et al., 2014). In eCLIP data (for is lacking for MATR3), ~10% of PTBP1 crosslink events mapped to antisense L1 elements, compared with ~4% across all RBPs examined. The decreased enrichment in eCLIP compared to iCLIP likely stems from differences in data processing (see Methods for comment).

Within our set of iCLIP data, we also found CELF2, ELAVL1 and TARDBP overrepresented at antisense L1s. Nine additional RBPs showed enrichment on L1 elements in eCLIP data: SUGP2, hnRNPM, KHSRP, hnRNPC, FUBP3 and SFPQ on antisense L1 repeats, and HLTF, KHDRBS1 and SAFB2 on L1 elements in sense. We also examined RBP binding to L2 elements, which are about three times less common in human genome than L1s. Correspondingly, we found that L2s accounted for a smaller proportion of RBP binding sites than L1s. SUGP2, MATR3, PTBP1 and HNRNPK had strongest enrichment in sense L2s, and HNRNPA1, TAF15, HNRNPU and SAFB2 in antisense L2s (Suppl. Table 2).

In our analysis we used reads mapping uniquely to the genome, which could miss the reads mapping to highly repetitive sequences. To account for them, we also examined eCLIP RBP binding to different sub-families of LINEs by using the TEtranscripts method (Jin et al., 2015). Median enrichment of LINE subfamilies recapitulated our ranking, with equal enrichment for all of the subfamilies for most of the RBPs and with strongest variation between families seen for HNRNPM and SFPQ (Figure S1A). In total, TEtranscript identified >2-fold enrichment on L1 or L2s for 25 RBPs in the eCLIP data. In conclusion, we find strong enrichment of MATR3 and PTBP1 binding on L1 and L2 elements, which appears to be related to their deep intronic binding profiles.

### MATR3 stabilises PTBP1-RNA interactions to promote L1 binding

MATR3 and PTBP1 directly interact (Coelho et al., 2015), but it is not known if this affects their endogenous RNA binding. Given that both proteins are enriched on antisense L1s, we wished to understand if their binding sites overlap. Since MATR3 eCLIP data are not yet available, we focused solely on iCLIP analysis. We analysed the five RBPs with most LINE binding in iCLIP and performed unsupervised clustering on the 50 most strongly bound LINEs of each RBP. The strongest correlation on individual LINEs was observed between MATR3 and PTBP1 (pearson coefficient = 0.83), with less overlap shared with CELF2, essentially no overlap with TARDBP-bound elements, and a slight anti-correlation with ELAVL1 occupied loci (Figure S2A). Conversely, MATR3 binding was enriched in proximity of PTBP1 binding peaks and this enrichment is significantly stronger at peaks located within rather than outside of LINEs (p-value < 2.2e-16, Figure S2B). Hence, LINEs appear to act as a platform for simultaneous binding of MATR3 and PTBP1.

Next, we examined if MATR3 and PTBP1 affect each other in their binding to LINEs. We performed iCLIP with PTBP1 in HEK293 cells depleted of MATR3, and iCLIP with MATR3 in HEK293 cells depleted of PTBP1 and PTBP2, and in both cases performed control iCLIP from cells transfected with control siRNA (Figure 2A and Figure S2C-D). Efficient siRNA-depletion of MATR3 and PTBP1 was validated by Western blot (Figure 2A and Figure S3D). Notably, the amount of RNA crosslinked to PTBP1 was visibly decreased upon MATR3 depletion, as measured by ^32^P labelling of immunoprecipiated RNA (Figure 2A; replicates in Figure S2C). This decrease was not a result of decreased abundance of PTBP1 (Figure 2A). We did not observe any decrease in MATR3 crosslinked RNA upon depletion of PTBP1/PTBP2 (Figure S2D). This indicates that MATR3 is required for efficient crosslinking of PTBP1 to endogenous transcripts, but not vice versa.

**Figure 2:**
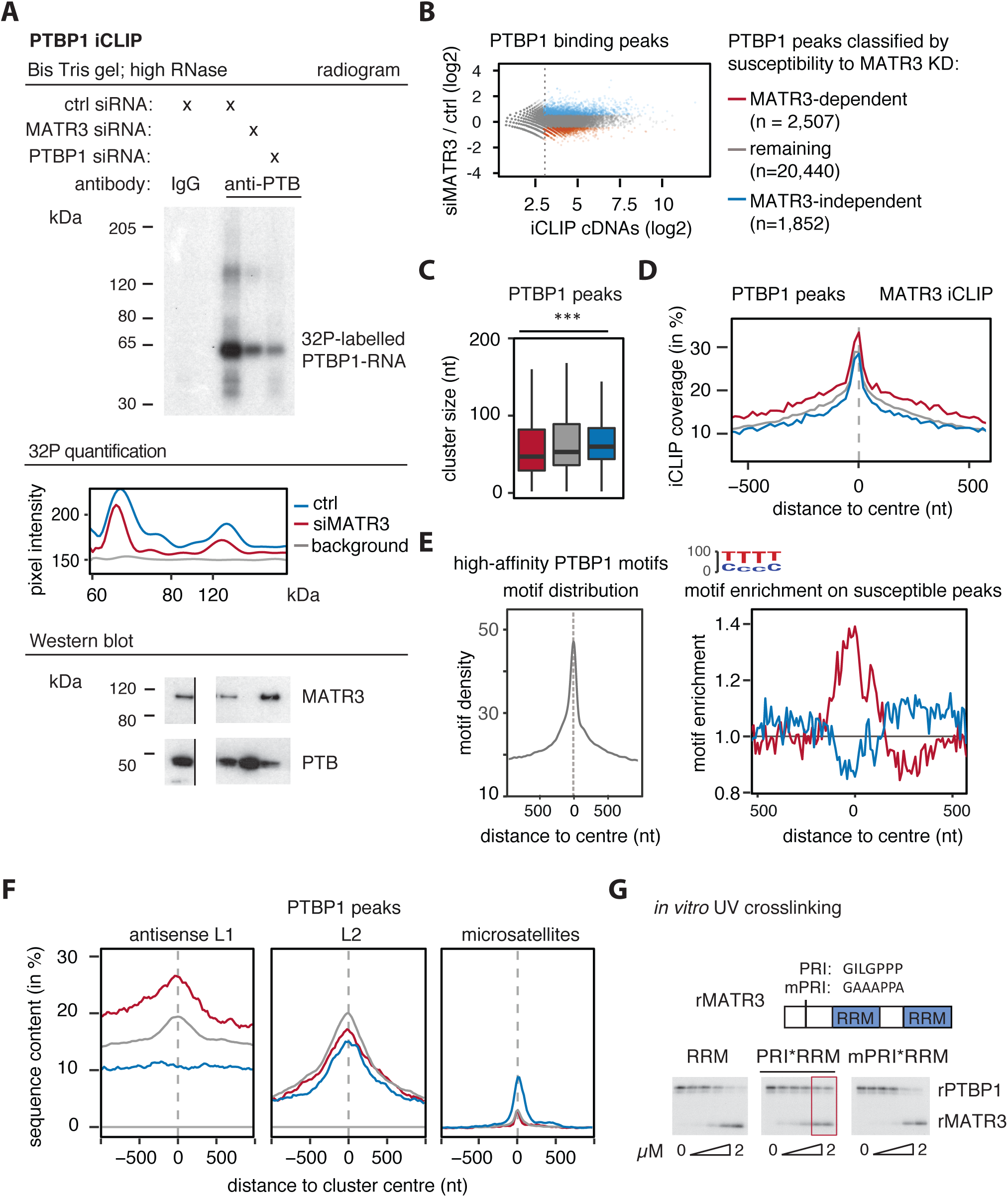
Binding of PTBP1 to antisense L1 elements is MATR3-dependent. PTBP1 iCLIP was performed from HEK293T cells depleted of MATR3, PTBP1 as well as controls. (A) TOP: ^32^P labelled RNA crosslinked to and co-precipitated with PTBP1 under high RNase conditions. MIDDLE: To quantify the signal, grey pixel intensity if shown across the centre of each lane, analysed with ImageJ software. BOTTOM: The input lysate for the iCLIP experiment was probed for MATR3 and PTBP1 antibodies in a Western Blot to ensure reduced signal is not due to changes in protein abundance. Samples are the same as in the radiogram, but the gel image was cut to align them. Note replicates are shown in Fig. S2A. (B) PTBP1 binding peaks were identified from all iCLIP experiments, and classified according to susceptibility to MATR3 depletion as indicated based on moderated log2 fold changes. Binding peaks with a normalised count of less than 8 were ignored, indicated by the dotted line. (C) PTBP1 binding peaks susceptible to MATR3 depletion are shorter than those which are not. (D) MATR3 iCLIP is enriched around MATR3-dependent PTBP1 binding peaks. (E) Enrichment for high-affinity motifs around PTBP1 binding peaks. LEFT: all PTBP1 binding peaks show strong enrichment for PTBP binding motifs. RIGHT: MATR3-dependent PTBP1 binding peaks show enrichment in a 200nt region for high-affinity motifs above other PTBP1 binding peaks. (F) The overlap between the centre of PTBP1 binding peaks and different repeat classes was tested for antisense L1 elements, sense L2 elements, and sense CT-/T-rich microsatellite repeats. Metaprofile shows percent of each class of clusters overlapping with each genomic element. MATR3-dependent binding peaks are more frequently derived from an antisense L1 element than MATR3-independent once. (G) Protein-protein interaction between MATR3 and PTBP1 allows formation of a heteromeric complex on a substrate RNA with two ATGTT motifs *in vitro.* Recombinant PTBP1 (rPTBP1) and different MATR3 mutants (rMATR3) were crosslinked to the same RNA at different MATR3 molarity (rPTBP1 at 0.5μM).

To analyse changes in PTBP1 binding upon MATR3 depletion, we identified peaks of PTBP1 crosslinking, focused on those with sufficient coverage, and classified them based on the change in normalised counts between MATR3-depleted sample and control into MATR3-dependent, MATR3-independent and remaining peaks (Figure 2B). MATR3-dependent PTBP1 peaks were shorter than MATR3-independent ones, and had more MATR3 binding in their vicinity (Figure 2C and D). As expected, PTBP1 binding peaks were highly enriched for CT-tetramers, which is most prominent within the peak, but is also seen over a 200nt region around the peak (Figure 2E). Intriguingly, we found that the enrichment over this 200nt region is stronger at the MATR3-dependent PTBP1 peaks compared to the remaining peaks (Figure 2E). Thus, the MATR3-dependent PTBP1 binding peaks are shorter but are composed of the highest density of binding motifs over a 200nt region. This indicates that MATR3 supports the interactions between PTBP1 and multivalent RNA binding sites, i.e., those that contain multiple repeats of high-affinity PTBP1 binding motifs

Next, we examined enrichment of different categories of peaks within repetitive elements. Importantly, MATR3-dependent PTBP1 binding peaks were most enriched within an antisense L1 elements compared to the remaining peaks (Figure 2F). PTBP1 also binds CT- and T-rich microsatellite repeats (Ling et al., 2016), but this accounts for only ~0.2% of all binding peaks in unperturbed HEK293 cells. While no L1 enrichment is seen for MATR3-independent PTBP1 peaks, they more frequently overlap with the microsatellite repeats. In the reciprocal analysis of MATR3 iCLIP after PTBP1 depletion (Figure S2D-F), we found that PTBP1 is not required for MATR3 binding to LINE repeats, indicating that MATR3 is recruited to LINEs either through its own specificity, or through interactions with additional RBPs.

Our analysis suggested that *in vivo* binding of PTBP1 to L1 repeats is stabilised by MATR3. To confirm that MATR3 directly aids RNA-binding of PTBP1, we purified recombinant PTBP1 (rPTBP1) and MATR3 fragments (rMATR3) that do or do not interact with PTBP1. We previously found a <u>P</u>TBP1 R<u>RM2</u> <u>Interacting</u> (PRI) motif N-terminal of the MATR3 RRMs is essential for interaction with PTBP1 RRM2 (Coelho et al., 2015). We purified a MATR3 fragment comprising its two RRMs but missing the PRI motif (‘RRMs’), as well as the RRMs with the PRI motif (‘PRI-RRMs’) and a mutated sequence with point mutations in the PRI disrupting PTBP1 binding (‘mPRI-RRMs’). We designed an *in vitro* synthesised RNA with two MATR3 RNA compete motifs (ATCTT, Ray et al., 2013) as well as small CT-stretches, which could allow multivalent binding of PTBP1 in vicinity. In agreement, rPTBP1 and all rMATR3 fragments bound to this RNA. We found that the non-interacting rMATR3 RRMs fragment competes with PTBP1 for RNA binding at equimolar concentrations (Figure 2G). Unlike the RRM rMATR3 fragment, the PRI-RRM rMATR3 did not block crosslinking of PTBP1 to the RNA even at excess molarity of rMATR3. The mPRI-RRMs rMATR3 blocked PTBP1 binding, demonstrating the dependency on the interaction motif for formation of a heteromeric PTBP1*MATR3 complex on the RNA. As a next step, we added rMATR3 to HeLa nuclear extracts with endogenous PTBP1, and assayed binding to two RNA probes. We used the probe with two ATCTT motifs (as in Figure 2G), and in addition a probe with six CTCTT motifs (the RNA compete motif for PTBP1), for which we expected stronger binding of PTBP1. Addition of rMATR3 promoted binding of endogenous PTBP1 to the exogenous ATCTT_2_ RNA through the PRI motif (Figure S2G). On the CTCTT_6_ probe, we observed increased binding of endogenous PTBP1 in absence of recombinant MATR3 compared to the ATCTT_2_ probe, and PTBP1 was completely displaced by non-interacting rMATR3 RRMs but not by the PRI-RRM rMATR3. Hence, the PRI motif in MATR3 allows formation of a heteromeric complex on substrate RNAs.

Together, we show that PTBP1 and MATR3 *in vivo* overlap at antisense L1-derived binding sites, which are rich in multiple repeats of high-affinity PTBP1 binding motifs. We find that the PRI-mediated interaction between MATR3 and PTBP1 is crucial to promote simultaneous binding of both proteins to an RNA, and that MATR3 can recruit PTBP1 to RNA in a sequence-dependent manner *in vitro* and *in vivo.* We conclude MATR3 promotes binding of PTBP1 to multivalent binding sites within antisense L1 repeats.

### MATR3 and PTBP1 co-repress exons and poly(A) sites close to LINE repeats

To resolve the role that coordinated LINE binding of MATR3 and PTBP1 might play in RNA-processing, we first re-analysed our previous splice junction microarray data on repression of alternatively spliced exons by MATR3 (Coelho et al., 2015). Of the 421 exons that were found to be repressed by MATR3, 64 contained at least one of their splice sites within a LINE repeat, and were therefore considered ‘LINE-derived exons’; this represents a 2.3-fold enrichment for LINE-derived exons compared to all exons covered in the array design. For PTBP1, we found 50 significantly repressed LINE-derived exons. We evaluated the frequency of L1 and L2 repeats in the introns flanking MATR3/PTBP1 repressed exons (Figure 3A, and Figure S3A), and found ~2-fold enrichment for antisense L1 sequence in a window of 2kb, even after removing all LINE-derived exons (Figure S3B). Hence, we found both LINE-derived as well as LINE-proximal exons are overrepresented among exons repressed by MATR3/PTBP1.

**Figure 3:**
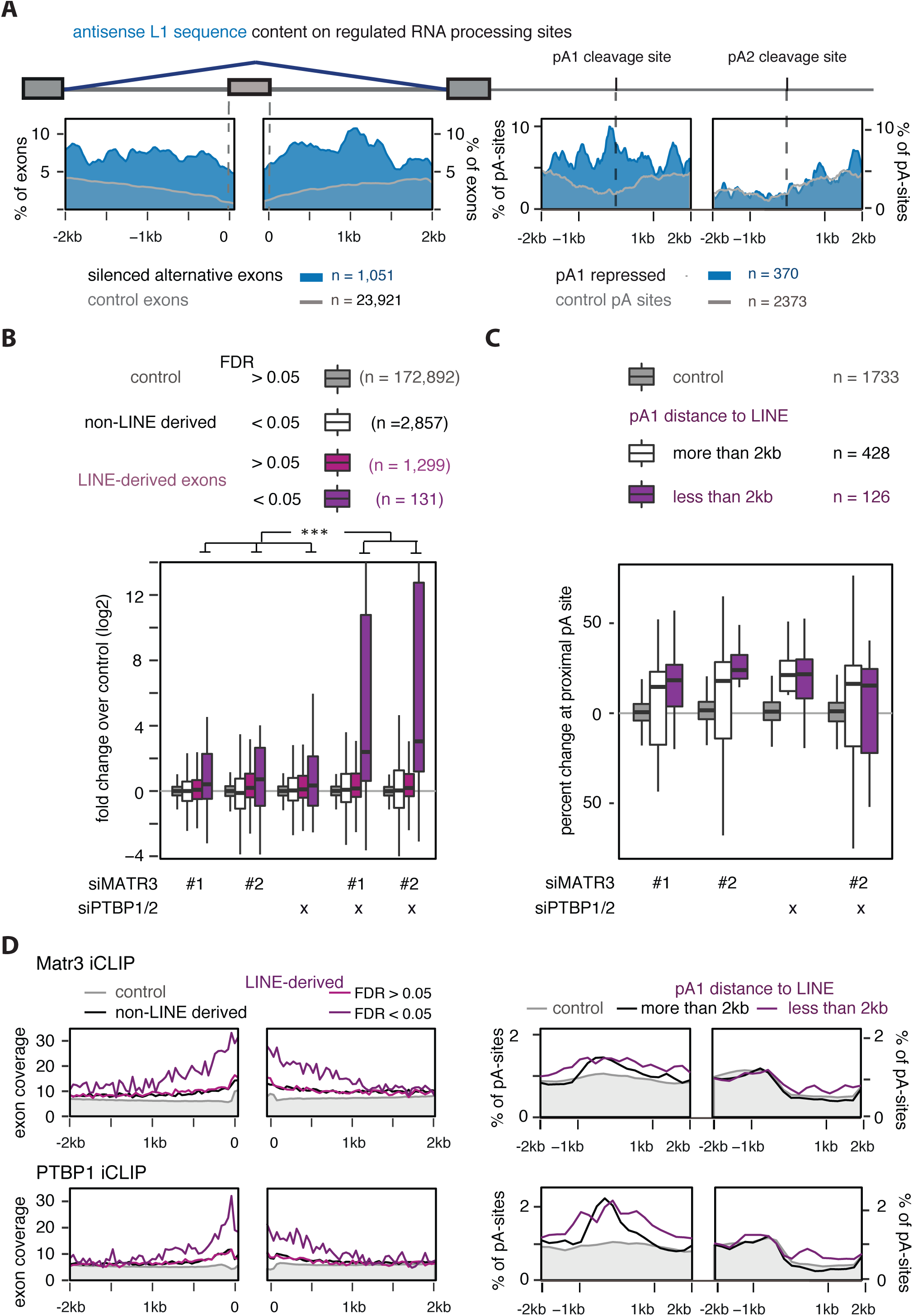
MATR3 and PTBP1 repress usage of cryptic splice and polyA-sites in vicinity to LINEs. (A) The metaprofile shows the coverage of antisense L1 sequences in a ±2kb window flanking the splice sites and the proximal and distal polyA sites of MATR3/PTBP1/2 repressed events. Exon usage and polyA-site usage was analysed in cells depleted of MATR3 and PTBP1/2, individually or in combination, and events significantly increased in absence of either proteins were selected. Misregulated exons are alternative exons selected from a splice-array experiment (Coelho et al., 2015), polyA site pairs are from mRNA 3’end sequencing experiments. Controls are non-significant events site with no appreciable change (below 10%) and reflect the expected genomic frequency of L1 antisense sequence (shown in grey). Metaprofile was smoothed using 40 nucleotide bins. (B) The transcriptome was *de novo* assembled from cells depleted of MATR3 and PTBP1/2, individually or in combination, in order to capture cryptic LINE-derived exons absent from microarrays. For each condition, the log2 fold changes of MATR3/PTBP1 regulated exons are plotted. Only events with at least one junction-spanning read were considered for analysis, and significant and non-significant LINE-derived exons are shown separately (at FDR < 0.05). Differences between the changes in exon abundance across groups were tested by Kruskal-Wallis Rank Sum test (p-value < 2.2e^−16^), and pairwise comparisons within each condition were tested with a two-sided Wilcoxon Rank Sum test, and corrected for multiple testing according to Bonferroni. Adjusted p-value indicated by *** was below 0.0001. Whiskers are cut-off from the boxplot for visualisation, but data distribution extends to (ymax +24) in cells depleted of MATR3 and PTBP1/2 simultaneously. (C) Percent change in usage of the proximal polyA-sites (same as in A). Misregulated pA-sites are split into those within 2kb vicinity of a LINE and those which are not. (D) Metaprofile of MATR3 and PTBP1 iCLIP binding across the splice and polyA sites ±2kb of the regulated event. Events were selected and grouped as in (B) and (C). iCLIP binding is presented as percentage of occupancy, and was smoothed using 40 nucleotide bins. Occupancy on non-regulated sites is shown as control (in grey).

Next we generated total RNAseq data of cytoplasmic and nuclear RNA from HeLa cells depleted of MATR3 with two independent siRNAs, or PTBP1/PTBP2, or all three factors simultaneously (Figure S3C and Suppl. Table 3). We detected 1,430 LINE-derived exons, each supported by at least one splice-junction read mapping to a LINE element; 1,114 within protein-coding genes and the remaining in long non-coding RNAs. Of the 1,430 exons, 858 (~77%) were cryptic, i.e. not annotated in UCSC (Suppl Table 2). LINE sequences can donate either 5’ or 3’ splice sites, and in ~50% of LINE-derived exons both splice sites were LINE-derived (Figure S3D). Depletion of both MATR3 and PTBP1 led to increased use of 131 (9.1%) of the LINE-derived exons (Figure 3B), with a median increase of more than five fold. Repression of these exons by the two proteins is strongly synergistic, since exon usage increased by about 1.6fold depleting MATR3 or PTBP1 individually. We tested changes in inclusion of 16 splicing events significantly regulated by co-depletion of MATR3/PTBP1 by semi-quantitative RT-PCR, including six cryptic and ten annotated exons (Figure S3E), and found synergistic repression for two out of nine LINE-derived exons and two out of five LINE-proximal exons.

Since antisense L1 elements are rich in cryptic polyA-signals (Han et al., 2004, Lee et al., 2008), we also produced 3’ end sequencing data to investigate if MATR3 and PTBP1 repress poly(A) sites in a LINE-dependent fashion. We used the expressRNA platform (Rot et al., 2017) to find alternative poly(A) site usage. We thereby annotated poly(A) site pairs in 5,189 genes, in which two different polyA sites each account for at least 5% of this gene’s signal (referred to as pA1 and pA2). Of these, 240 pA-sites originated from a LINE repeat. LINE repeats were enriched at proximal polyA sites repressed by MATR3/PTBP1 for an extended region of ~2kb (Figure 3A), reminiscent of the pattern observed on repressed exons. Overall changes in polyA site usage suggest a primarily repressive function of MATR3/PTBP1 (Figure 3C). We split all significantly regulated proximal pA-sites into those within 2kb of a LINE, and those further away from any LINE repeat, and found LINE-proximal sites to be slightly more responsive to MATR3 depletion than LINE-distant sites (Figure 3C). This is mirrored in individual examples (Figure S3 E; for instance in MROH1, an annotated alternative terminal exon with a LINE-derived 3’ SS (indicated by red dashed line) is used ~70% in control cells, but entirely replaces the canonical pA site in MATR3 depleted cells accompanied by exonisation of additional sequences and an additional pA site, all from the adjacent LINEs. In PIGN1 (Figure S3F), a stretch of LINE sequences give rise to a cryptic LINE-derived terminal exon, which is used partially upon MATR3 depletion, and fully upon combined depletion of MATR3 and PTBP1, as indicated by loss of all signal on the downstream exon.

Metaprofiles of iCLIP binding on regulated splice- and polyA-sites showed increased binding of MATR3 and PTBP1, confirming direct targeting of these loci. LINE-derived exons were enriched in MATR3 and PTBP1 binding compared to non-repeat derived exons (Figure 3D) and those LINE-derived exons most susceptible to depletion of MATR3/PTBP1 showed strongest enrichment. MATR3 binding was extended for ~2kb upstream to ~1kb downstream of the exons, covering both splice sites. At polyA sites, MATR3 and PTBP1 binding was enriched at repressed pA1 sites, with extended binding on those pA sites that were proximal to a L1 repeat.

We conclude that MATR3/PTBP1 are potent repressors of RNA processing at LINE repeats, thus preventing exonisation of LINEs. Similarly, polyA sites are repressed in vicinity of LINE repeats. Together, this strongly suggests that LINEs are the specificity element in directing MATR3 to alternative exons, linking its function in alternative splicing to its binding on repeat elements, and explaining the lack of a short binding motif of MATR3 *in vivo* we described in the past (Coelho et al., 2015). The binding pattern of PTBP1 on LINE-derived exons was consistent with co-targeting of the same elements by MATR3 and PTBP1. Lastly, changes in abundance of LINE-derived exons suggest functional synergy of MATR3 and PTBP1 on LINE-derived exons but not on non-repeat derived alternative exons, suggesting co-ordinated assembly of both proteins is necessary to ensure complete repression of cryptic exons originating from LINE repeats.

### LINE-derived exons reduce transcript abundance through NMD

The majority of LINE-derived exons that were detected in MATR3/PTBP1 depleted cells are cryptic exons, i.e. not annotated by UCSC or ENSEMBL (Suppl Table 2). Retrotransposon-derived exons, and in particular Alu-exons, are known to be prone to spurious inclusion which generally reduces expression of the host gene through nonsense-mediated mRNA decay (NMD) (Attig et al., 2016). Given the involvement of PTBP1 in repression of LINE-derived exons, we used RNAseq data produced by Ge et al. (Ge et al., 2016) to evaluate if LINE-derived exons detected in HEK293 cells trigger NMD. Depletion of PTBP1 alone produced a marked change in abundance of LINE-derived exons, while depletion of UPF1 alone drastically increased the number of LINE-derived exons detected (Figure S4A); and this number almost doubled after combined depletion of UPF1 and PTBP1. Importantly, genes containing any of those LINE-derived exons showed increased expression in UPF1-depleted cells (Figure S4B). We conclude most LINE-derived exons are cryptic exons that, when included, render the resulting transcript susceptible to NMD.

### Deletion of an intronic LINE disrupts MATR3-dependent repression of a cryptic exon in ACAD9

To confirm MATR3 and PTPB1 directly repress exons flanked by a LINE within 2kb of their splice site, even if they are not LINE-derived exons, we made a splice reporter plasmid. Among 16 exons for which we validated the role of MATR3 and PTBP1 by RT-PCR (Figure S3E), 6 exons were such LINE-proximal exons, including an exon within intron1 of ACAD9. Endogenous ACAD9 splices efficiently at intron1, with two known splice isoforms of exon1 using different 5’ splice sites (here referred to as *exon 1a* and *exon 1b).* We observed a two-fold loss of ACAD9 expression in cells depleted of MATR3, and a three-fold loss of expression in cells co-depleted of MATR3 and PTBP1 (Figure S4 C and D). Intron1 of ACAD9 contains three L2 repeat elements in sense orientation, which all showed pronounced binding by MATR3 and PTBP1 in cultured human cells as well as binding by MATR3 and PTBP2 in mouse brain (Figure S4C). We confirmed by RT-PCR that individual depletion of MATR3, but not PTBP1, led to inclusion of an alternative exon starting 323 bp upstream of the L2 repeats (Figure 4B), and verified its splice sites by Sanger sequencing. Notably, inclusion of the exon is markedly elevated after co-depletion of MATR3 and PTBP1 (Figure 4B).

**Figure 4:**
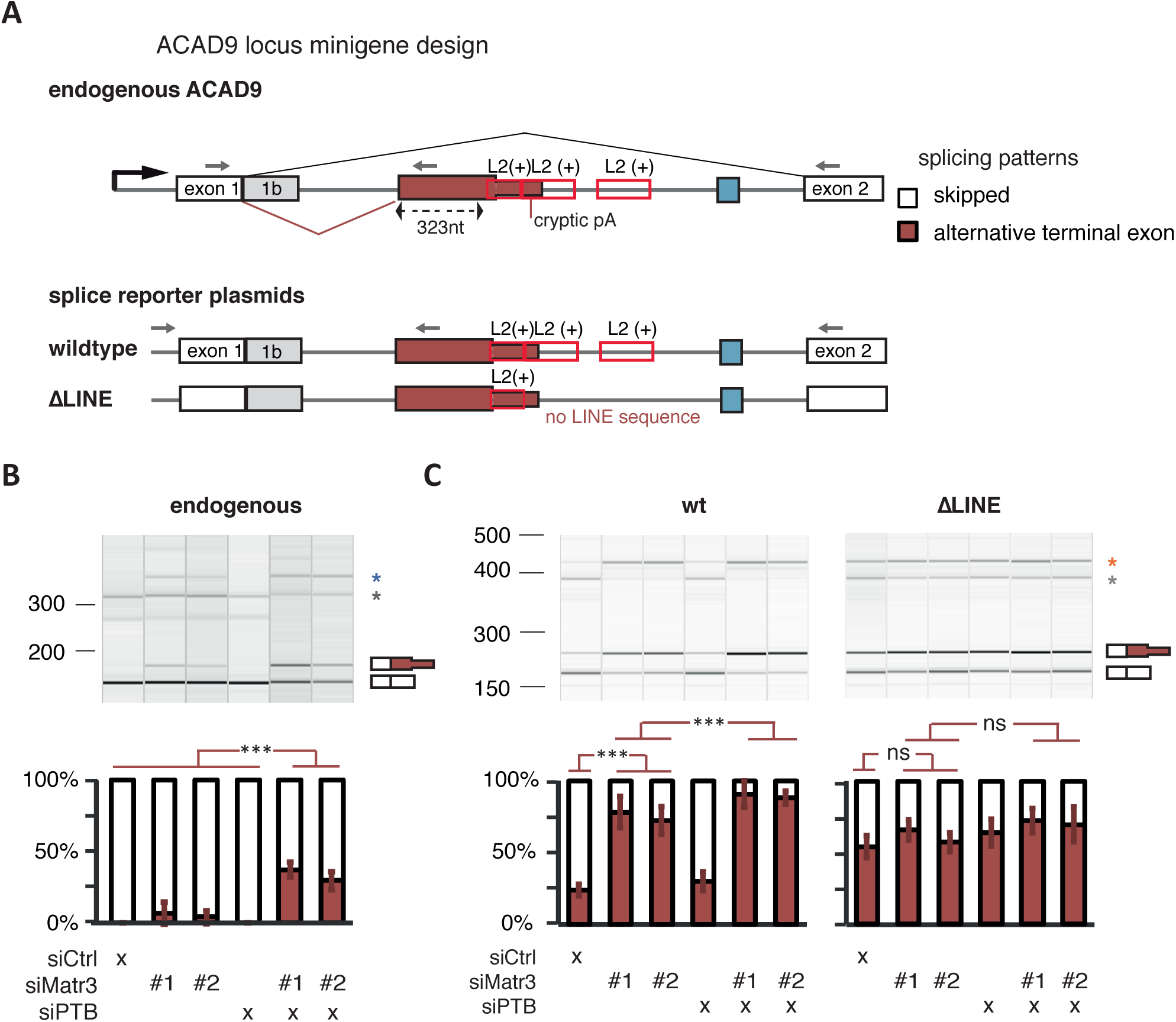
Partial deletion of L2 sequences disrupts splicing repression of *ACAD9* by MATR3/PTBP1. (A) Schematic illustrating the endogenous ACAD9 locus and the ACAD9 splice reporter. The first two exons and the complete intron1 were cloned into a CMV driven reporter plasmid. In the ΔLINE splice reporter 499 base pairs of L2 sequence were replaced by non-repetitive sequence of intron2 of ACAD9. (B) The inclusion level of the LINE-proximal alternative terminal exon in endogenous ACAD9 was measured in total RNA of cells depleted of MATR3 and PTBP1/2 individually or in combination as well as controls. To test for significance, one-way ANOVA was used coupled with multiple comparison correction according to Tukey’s HSD. *** indicates p-value below 0.001. Semi-quantitative RT-PCR analysis is averaged across three independent replicates, error bars represent s.d.m. (C) The inclusion level of the LINE-derived exon was measured as in (B) in the wild-type and ΔLINE ACAD9 splice reporter. (B) and (C) Additional splice products are indicated by asterisks. These use the 5’ splice site of exon 1b.

To confirm that the LINE nucleotide sequence recruits MATR3 and PTBP1 and causes distant splicing repression, we created a splice reporter of ACAD9 comprising exon1 and exon2 and the complete intronic sequence including all three L2 repeats (called wildtype), and a mutant splice reporter that lacked two out of the three L2 repeats (called ΔLINE, see Figure 4A). The wildtype reporter reproduced the splicing pattern of the endogenous sequence in non-perturbed cells and in cells depleted of MATR3, PTBP1 or both (Figure 4C), except of overall more frequent usage of the 5’ splice site of exon 1b. Importantly, the ΔLINE splice reporter showed increased usage of the LINE-proximal 3’ splice site in unperturbed cells, with little to no further change in incusion upon MATR3/PTBP1 depletion (Figure 4C). Hence, the L2 sequence downstream of the exon was essential to confer responsiveness to MATR3/PTBP1. This supports our transcriptome-wide finding that MATR3 and PTBP1 repress LINE-proximal exons, in addition to regulating LINE-derived exons.

### PTBP2 prevents LINE-exon inclusion in mouse brain

Having identified the role of MATR3 and PTBP1 in repressing the splicing of cryptic LINE-derived exons, we sought to explore if they might play such a role also in the brain, given the known role of the PTBP1 orthologue PTBP2 in regulating splicing during neuronal development (Li et al., 2014). We first generated iCLIP data of MATR3 from mouse brain, and compared the enrichment on LINEs in the mouse brain for PTBP2, MATR3, CELF4, FUS and TARDBP. Enrichment was most pronounced for MATR3 and highest on antisense L1 sequences, to a similar extent as in HEK293 cells (Figure S5A). Interestingly, we found MATR3 and PTBP1 show stronger enrichment on rodent-specific L1 families than on evolutionary older L1 families. A MATR3 knockout mouse is not available (MGI:1298379); therefore we focused on RNAseq data from PTBP2^−/−^ mouse brain (Li et al., 2014, Vuong et al., 2016). In nestin-Cre-PTBP2^−/−^ E18 mouse brain, we found LINE-derived exons were more likely to be significantly deregulated than SINE-derived exons or non-repeat derived exons (Figure S5B; x^2^-test, p-value < 10^−5^) and were repressed by PTBP2 (Figure S5B; x^2^-test, p-value < 10^−5^). Hence, we suggest repression of LINE-derived exons is redundant between PTBP1 and PTBP2. Focusing on exons with inclusion levels above 10% (measured as percent spliced index, or PSI, see SupplTable3 and 5 of (Vuong et al., 2016)), PTBP2-regulated exons in Emx-Cre-PTBP2^−/−^ mouse brain include LINE-derived exon3 of *Fam124A*, exon5 of *Osblp9* and exon 2 of Ub3g1, all of which modify the protein sequence. These exons are absent or lowly included in E14 wildtype brains but their inclusion increases in the adult brain at P10 (Figure S5C to E). Out of 13 LINE-derived exons selected for this pattern, 8 are rodent-specific insertions that are not shared between mouse and human. This suggests that PTBP2 preferentially represses exons derived from evolutionarily young LINEs during brain development, and several of these exons become more highly included in the wildtype adult brain, thus gaining the potential to alter the species-specific neuronal transcriptome.

### Evolutionarily old LINEs are a major source of mammalian alternative exons

After observing exonisation of LINEs in mouse brain, we decided to survey the inclusion of LINE-derived exons in human tissues by using the extensive data available from the GTEx consortium (V6p data (Consortium, 2015)). We tested the percent inclusion of a total of 45,940 exons in RNAseq data from 51 human tissue types, covering all exons of the 4,566 genes that contain a known LINE- or Alu-derived exon. We detected 1,154 LINE-derived exons with at least 5% inclusion in at least one tissue. The LINE-derived exons showed higher degree of exonisation than Alu-derived exons, measured by maximum inclusion level (PSI) across tissues (Figure S6A), but showed a similarly high degree of tissue-specificity (Figure S6B). In contrast to well-established alternative exons, Alu- and LINE-derived exons are virtually never switch-like events (Figure S6B).

Since we observed enrichment of PTBP2-regulated exons derived from evolutionarily younger LINE families in mouse brain (Figure S5C to E), we also further explored the evolutionary history of human LINE-derived exons. For this purpose, we determined the evolutionary age of all human L1 and L2 repeats by cross-species comparison with two primate genomes (gorilla and rhesus macaque), two rodents (mouse and rat), and one each of the carnivore and laurasiatherian genomes (dog and cow, respectively), which were chosen due to the high quality of their genome assemblies. In this manner, we annotated LINEs as primate-specific, euarchontoglires-specific or mammalian-wide insertions (Figure 5A). We were able to categorise mammalian-wide insertions further by assigning if they were present in dog and cow or only one out of the two, which might indicate differences in selection pressure. We ignored elements for which the evolutionary history remained unclear, or which were present but largely sequence truncated in dog or cow. The number of substitutions of the elements from family consensus validated the average age in our annotation (Figure S6D), although we found it to be more robust on L1 than on L2 elements. Human L2 elements are much older than L1s (Deininger and Batzer, 2002), which means their insertion age is frequently older than the divergence of the genomes we used. Since it remains unclear if any L2 family has remained active in early ancestors of the euarchontoglires lineage, we focused on L1 elements for analysis of young LINE insertions.

**Figure 5:**
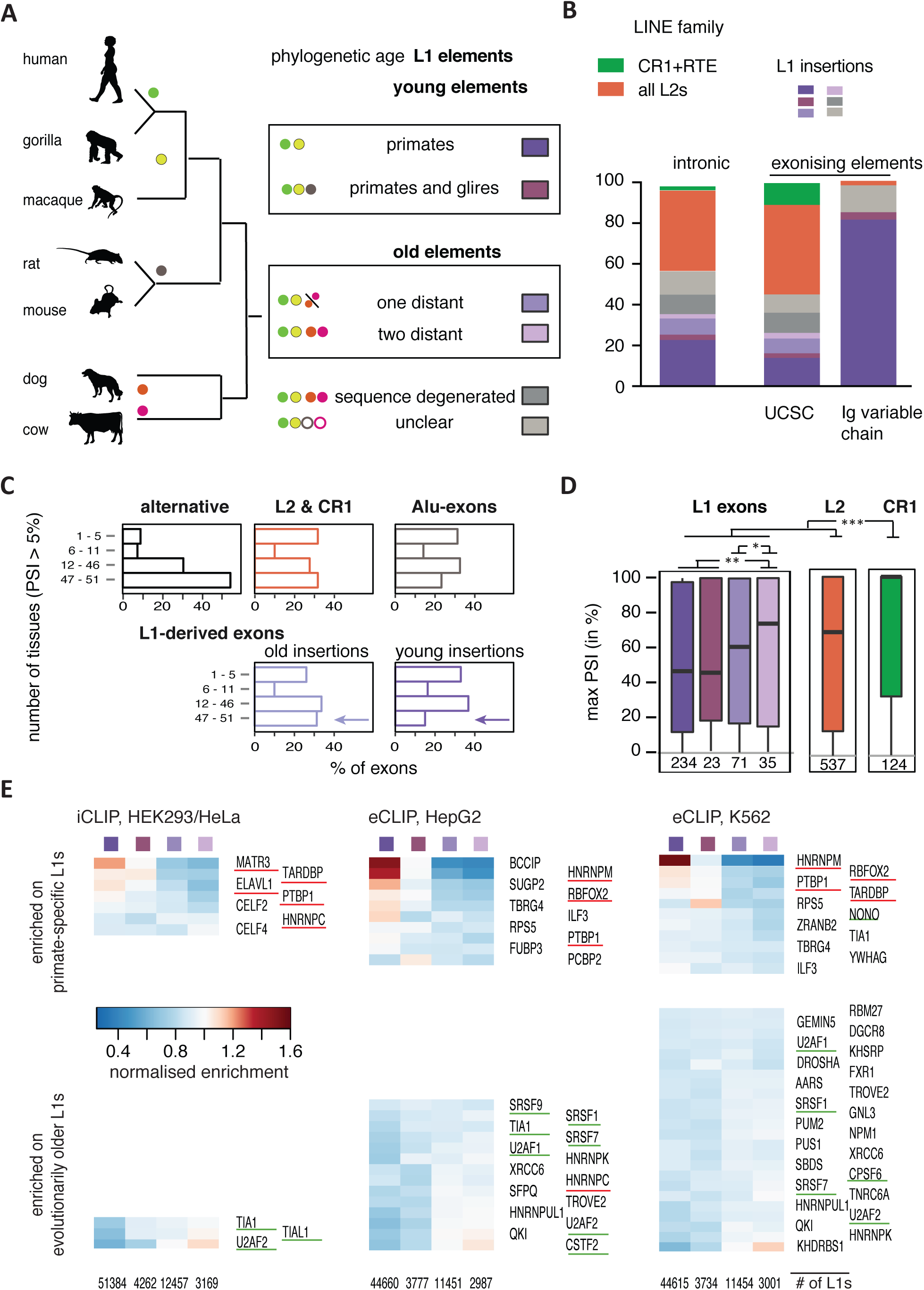
LINE-derived exons are a source of primate-specific alternative exons. Percent splice index (PSI) was calculated in the GTEx panel of human tissue samples for LINE-derived exons annotated in UCSC (relative to the flanking exons). Inclusion levels range from 0 to 100%, showing no inclusion or full inclusion. If no support for expression of the flanking exons was found, the gene was assumed to be non-expressed. (A) The phylogenetic age of each LINE element in the human genome was mapped by comparison to the gorilla, rhesus macaque, mouse, rat, dog and cow genome assemblies using UCSC liftover genome alignments overlaid with RepeatMasker annotation (see Methods for details). Elements specific to the primate or euarchontoglires lineage are considered evolutionarily young elements, while elements present in cow and dog are considered old elements. (B) The phylogenetic age of a LINE element gave an estimate of the genomic age of each LINE-derived exon. UCSC annotated exons are generally of the youngest elements. Within UCSC, the Ig-encoding region (*abParts*) stands out with 1,152 out of 6,012 annotated LINE-derived exons, which are frequently primate-specific. (C) Exons derived from evolutionarily young L1 elements are rarely present across human tissue subtypes. We determined the number of tissues in which each exon was detectable (at PSI > 5%) and compared repeat-derived exons to non-repeat derived known alternative exons. (D) Maximum inclusion in any tissue correlates with the genomic age of L1-derived exons. Significance was tested across groups by Kruskal-Wallis’ Rank Sum test and pairwise comparisons by Dunn’s test corrected according to Holm-Šidák. *, ** and *** indicate adjusted p-value was below 0.05, 0.01 and 0.001, respectively. (E) RBPs show preferences for binding to L1 elements of different evolutionary ages. The L1 elements with 10% highest coverage across any i/eCLIP data were used to calculate a normalised coverage for each RBP, and the number of L1 elements in each group is given at the bottom. Binding of each RBP was normalised by the sum of all RBPs within each cell line on an individual L1 element to obtain a relative binding estimate, and for visualisation of binding preference, normalised enrichment of each RBP was calculated by normalising to the mean.

Next, we examined the proportion of L1 repeats from each phylogenetic LINE group that is capable of seeding exons. We were surprised to find that L1-derived exons are a rich source of exons in the regions of the genome that encode the highly variable and species-specific immunoglobulin variable chain region (the Ig-region on chromosomes 2, 14, 15, 16 and chr22, Figure 5B). The Ig-regions are densely packed with 1,845 LINEs, 1,152 of which produce exons according to exon annotation by UCSC. The LINE-derived exons in these regions are almost exclusively seeded by primate-specific L1s (Figure 5B), and we consider them as cryptic exons, since we did not detect them by our analysis of the GTEx data. However, even when they are included, the exons are unlikely to map to the reference genome due to the rearrangement of the variable chain region during B- and T-Cell maturation. Detailed analysis of B and T cell receptor sequences will be needed to further examine the contribution of these young L1-derived exons to the expression of immunoglobulin genes. After excluding the Ig-regions, we find that less than 0.4% of the evolutionary young L1s can seed exons compared to 0.8% of the well-preserved older L1 elements, demonstrating that the older L1s more frequently contribute to the established transcriptomes across tissues (Figure S6C).

To quantify the differential regulation of LINE-derived exons across tissues, we calculated the maximum difference in inclusion between any pair of tissues. Interestingly, the inclusion of exons seeded by young L1 elements was more tissue-specific (Figure 5C, Figure S6E). However, exons derived from evolutionarily older LINEs generally showed higher maximum inclusion, comparing young L1 elements to either old L1 elements, or to exons derived from L2 and CR1 elements (Figure 5D). We found 594 L2- and 150 CR1-derived exons, which had inclusion levels similar to the evolutionary old L1 insertions (Figure 5D). In fact, CR1-derived exons were the group with highest inclusion levels of all groups examined, which agrees with them generally being the evolutionarily oldest in human. Between tissues, we found most LINE-derived exons in tissues of the reproductive system (Testis, Fallopian tube and Cervix) and the brain (considering all 13 regions of the brain; Figure S6G). Taken together, our analyses show that the evolutionarily older LINE insertions are a major source of mammal-specific alternative exons, some of which have reached high inclusion levels in different human tissues.

### Loss of repressor binding drives the exonisation of LINE-derived exons

To explain the differences in the inclusion level of the different evolutionary categories of LINE-derived exons, we compared their splice site strength, but did not find any marked differences (Figure S6F). Therefore, we reasoned that instead of changes in splice site strength, changes in the binding of different RBPs might determine exonisation of LINE-derived exons. To test this hypothesis, we exploited the available iCLIP and eCLIP data to analyse trends in RBP binding across the phylogenetic groups of L1 insertions. To ensure that we assessed elements that are part of expressed transcripts, we selected the 10% of L1 elements with highest coverage by any of the 121 RBPs. All phylogenetic groups were represented in this selection in expected proportions. Next, we averaged the binding of each RBP against the sum of all RBPs, generating a relative binding metric among all RBPs (ranging from 0 to 1). We then visualised any preferences in binding to a phylogenetic group as enrichment, considering all 49 RBPs that had above-average binding to LINEs (see Figure 1A). Strikingly, MATR3 is the RBP with strongest enrichment on primate-specific L1s among iCLIP experiments, and BCCIP and hnRNPM among eCLIP (Figure 5E). Both iCLIP and eCLIP, in both cell lines, also show PTBP1 enriched on primate-specific L1s. In general, known splicing repressors are enriched on primate-specific L1s, with the exception of hnRNPC. In contrast, RBPs that are well known to enhance splicing or 3’ processing also bind to evolutionarily older L1s, which includes SR proteins, and RBPs that recognise sequences close to 3’ and 5’ splice sites, or the polyA sites (Figure 5E, lower panels). We conclude that the stronger binding of repressive RBPs to the young L1s is the likely reason for their lower inclusion. The loss of these repressive RBPs, accompanied with binding by splice-promoting factors, can thus explain why the evolutionarily older L1s are the most common source of exons, and why they tend to be more highly included.

### Sequence divergence of the evolutionarily old L1 elements results in loss of repressor binding

To understand why the evolutionarily older LINEs do not bind repressive RBPs as well as younger insertions, we analysed the density of sequence motifs known to interact with these RBPs (Figure 6A). We found binding motifs in the literature for ELAVL1, HNRNPK, HNRNPM, KHDRBS1, QKI, PTBP1/2, RBFOX1 and TARDBP (see Suppl. Table 1 for details and references). We tested the distribution of all 256 tetramer sequences and found clear differences in line with the expected AT-richness of antisense L1 sequences (Suppl. Table 6), with TG- and TA-rich motifs being quite abundant in antisense L1 and CG-rich motifs being most depleted. We found evolutionarily older L1s contained fewer binding motifs of hnRNPM, TARDBP and ELAV1 (at FDR < 0.1). We found they contained on average more binding motifs of KHDRBS1, hnRNPC and QKI, though a large proportion of evolutionarily old L1s did not contain any QKI motif and none of the motifs associated with these three was among the most enriched motifs. PTBP1 motifs were highly abundant (a median of 1.26 motifs per 100nt) in all L1s, irrespective of their genomic age. We conclude the unequal binding towards L1s of RBPs, especially of splicing repressive RBPs such as hnRNPM, ELAVL1 and TARDBP, is a consequence of the L1 sequence and its decline through accumulation of sequence mutations.

**Figure 6:**
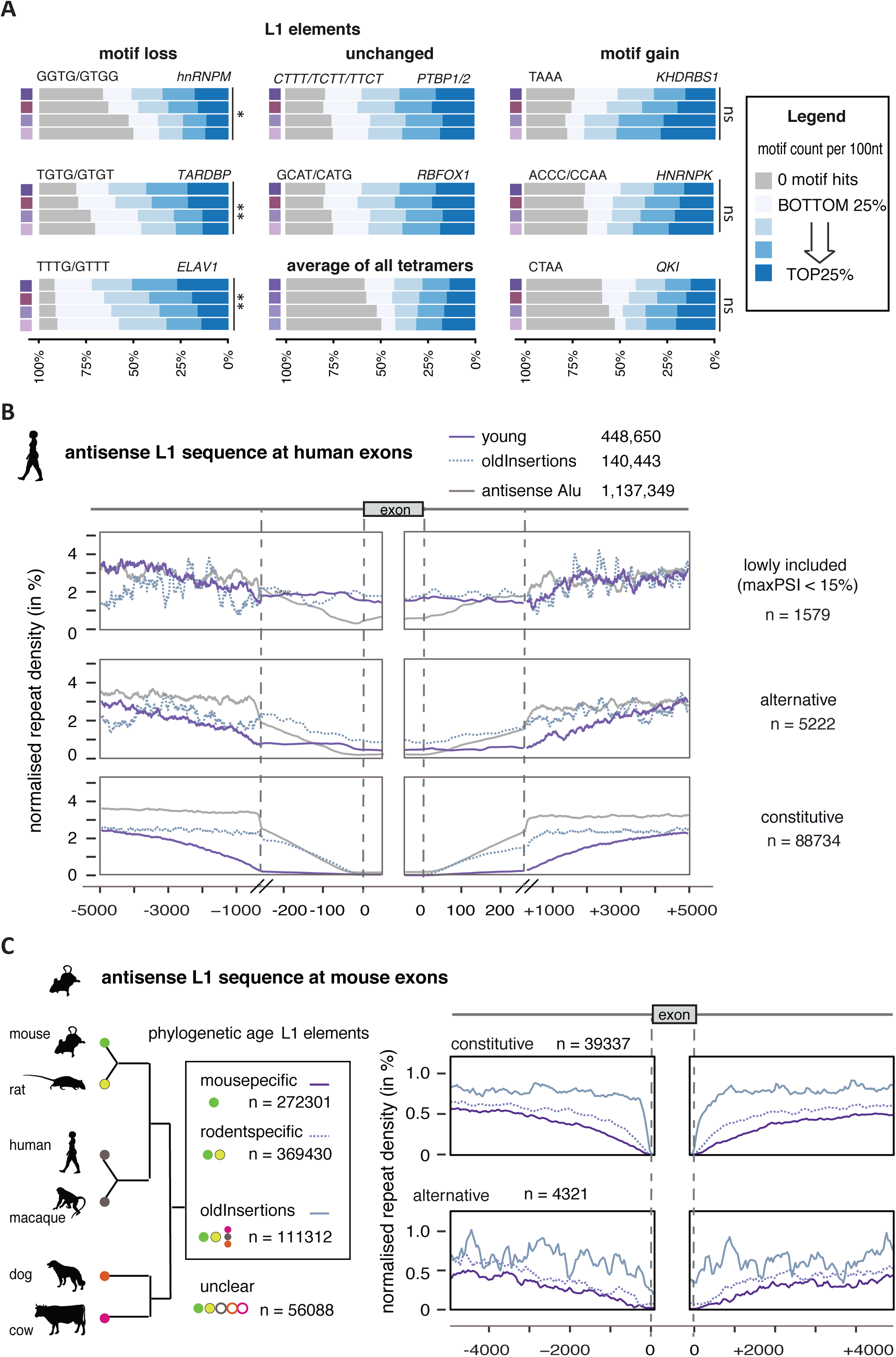
Young L1 elements are rich in splice repressor binding motifs and selected against at exons in a broad window. (A) The number of binding motifs associated with different RBPs is shown for antisense L1 sequences of different genomic age. Binding motifs of RBPs shown in Figure 5E were identified from literature where possible and searched for in antisense L1 elements. The genomic age of L1 elements is defined as in Figure 5A. Total motif count per 100nt was determined and categorised as quartiles (bottom to top 25% and 0 motifs, see legend). For comparison, the average distribution of all possible tetramers is shown. Changes in motifs counts with evolutionary age of the elements were considered significant based on their empirical distribution (see Methods for details)., ** and * indicates FDR below 0.05 and 0.1, respectively; ns = not significant. (B) Density profiles showing L1 antisense sequence 5kb upstream and downstream of human exons. L1s were split for evolutionary young and old insertions and repeat density is normalised to the total number of repeats in the two groups. For comparison, the primate-specific Alu insertions are shown. Exons were grouped by inclusion in human tissues from GTEx data into those which are more than 5% but less than 15% included in any tissue, those which are alternative and those which are constitutively included. To better present the repeat density around the splice sites, the x axis is cut at 250 nt to show a zoom-in of the 250nt flanking the exons. (C) Density profiles showing L1 antisense sequence 5kb upstream and downstream of constitutive and alternative exons in the mouse. The genomic age of each L1 element in the mouse genome was mapped by comparison to the rat, rhesus macaque, human, dog and cow genome assemblies (see Methods for details).

### The evolutionarily young LINEs maintain the repression of deep intronic regions

Given that we show assembly of mostly splice-repressive RBPs at and across evolutionary young LINE sequences, we hypothesised that these LINEs are in a repressed state. Furthermore, at least MATR3 and PTBP1 inhibited splicing also in nearby regions, which raised the intriguing question of whether evolutionarily young LINEs are generally prohibitive for splicing. To test if young L1s act as intronic splice silencers, we examined their distribution around annotated exons as well as the inclusion of these exons across human tissues. Strikingly, we found that evolutionarily young LINEs were excluded from an approximately 3kb region around constitutive and alternative exons (Figure 6B). However, they were not excluded around those exons with very low inclusion across human tissues (maxPSI<15%), indicating that they may contribute to the repression of these exons (Figure 6B). In total contrast to young antisense L1 sequences, the primate-specific Alu repeats were only excluded from the immediate vicinity of exons, but not from flanking intronic regions. Older L1 elements are well tolerated up to 250bp at all exons, and their incidence decreases only within ±200nt of constitutive exons. As independent validation, we repeated the analysis on mouse exons, and found mouse- and rodent-specific LINEs excluded in a large window around their splice sites, a pattern recapitulating the primate-specific insertions in human (Figure 6C). Thus, the evolutionarily younger antisense L1 elements are more depleted from the vicinity of exons both in primates and rodents. This could be a consequence of them being particularly potent in recruiting repressive RBPs such as MATR3 and PTBP1, which leads to repression of exons in their vicinity.

## DISCUSSION

We find that tens of thousands of LINEs recruit a diverse set of 28 RBPs to deep intronic loci. Insulating RNA from the splicing and polyadenylation machinery is a known mechanism of repression used by a number of RBPs (Witten and Ule, 2011). Of the RBPs binding of LINEs many are splicing repressors, such as MATR3 and PTBP1, which repress the recognition of LINE-derived exons and polyA sites, both in cultured human cells and in PTBP2^−/-^ mouse brain. Importantly, we demonstrate that MATR3 promotes efficient PTBP1 binding to L1s by stabilising its interaction with multivalent binding motifs.

The repetitive nature of LINEs and their evolutionary divergence allowed us to demonstrate a dual role of young and old LINEs in RNA processing (Figure 7). Repressive RBPs preferentially bind to the young LINEs and insulate both the LINEs and the surrounding regions from the processing machineries. As a consequence, young LINEs are confined to deep intronic regions. We propose their insulation allows the accumulation of cryptic RNA processing sites and facilitate evolutionary innovation. Through the accumulation of sequence mutations, the density of repressive binding motifs is gradually decreased, and these processing sites become gradually de-repressed. This is evident by the closer proximity of older LINEs to exons, by their binding to RBPs that enhance RNA processing, and by their increased contribution to tissue-specific transcript isoforms. Thus, we find that intronic LINEs play a dual role: they recruit repressors to insulate the deep intronic regions from processing machineries, but after long evolutionary periods also act as a source of RNA processing sites that facilitate the formation of mammal-specific transcripts.

**Figure 7:**
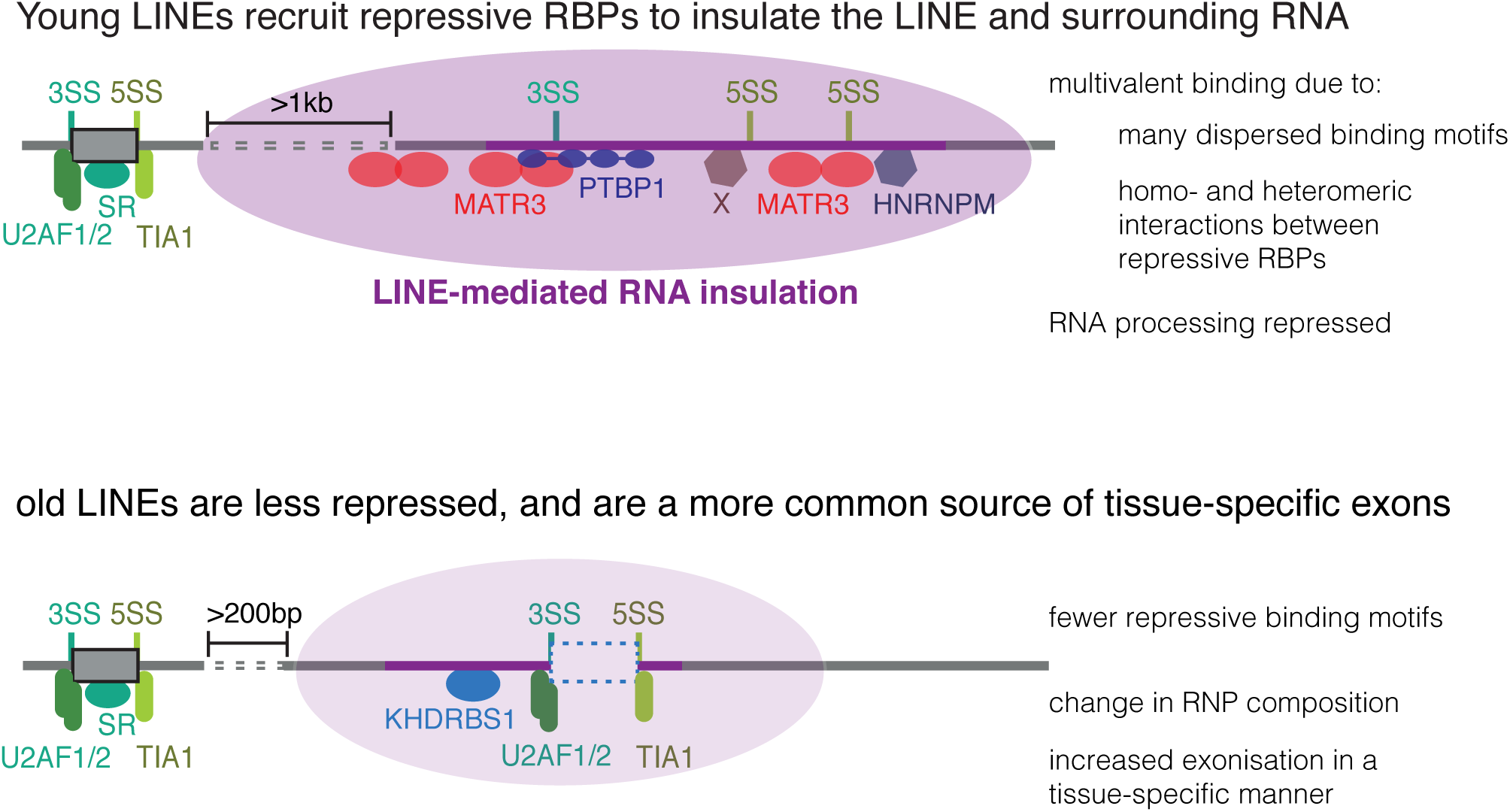
LINE elements create a splice repressive zone that prevents cryptic exonisation events. Consensus L1 elements are known to contain strong splice sites, but exonisation is rare and generally we observed exons from elements that are evolutionarily old. Evolutionarily young L1 insertions recruit a number of splice repressive proteins, including MATR3, PTBP1 and hnRNPM, as well as RBPs of yet unknown function (indicated as an X; but candidates are for instance BCCIP and SUGP2, see Figure 5C). These proteins recognise RNA motifs present within the L1 elements, which are diminished within evolutionary older L1s. The extent of splice-repressive proteins assembling on the L1s leads to selective pressure against young L1 insertions in a large proximity window of non-repeat derived exons. Hence evolutionary young LINEs insulate intronic regions from RNA processing. Evolutionarily older elements have a high probability of loosing binding sites of repressive RBPs. Hence, their exonisation is more common, but still largely tissue-specific.

### MATR3 and PTBP1 bind LINEs to synergistically repress RNA processing

We found that antisense L1 and sense L2 elements recruit MATR3 and PTBP1 to deep intronic regions, where both RBPs repress RNA processing at and around the bound LINEs. Binding of both proteins significantly overlaps at these LINEs, and MATR3 is required for efficient PTBP1 binding to L1s. Antisense L1s contain many PTBP1 binding motifs and MATR3-dependent binding sites of PTBP1 are characterised by increased density of binding motifs over a broad 200nt region. A model to explain our observations is that LINEs provide multivalent binding sides, and complex formation with MATR3 promotes PTBP1 binding to those. PTBP1 is capable of multivalent RNA binding through its four RRM domains (Oberstrass et al., 2005). Analysis of liquid phase separation properties of PTBP1 *in vitro* recently demonstrated that its binding is mediated in part by multivalent binding sites on the RNA, and is further stabilised by fusing PTBP1 to intrinsically disordered regions (IDRs) of different proteins, due to IDR-mediated protein-protein interactions (Lin et al., 2015). It was therefore proposed that the PTB-RNA and IDR interactions could act together to produce larger oligomeric assemblies with increased affinity for RNA. PTBP1 RRM2 interacts with a short linear motif within an IDR in MATR3 (the PRI motif, Coelho et al., 2015), and MATR3 interacts with itself (Zeitz et al., 2009). It seems likely that PTBP1 and MATR3 assemble across the antisense L1 sequences as a larger oligomeric assembly through multivalent RNA binding. Notably, we find that one of the previously studied MATR3-repressed exons is derived from a sense L2 insertion (exon 11 in ST7), and repression of this exon depends on its PRI motif (Coelho et al., 2015), indicating that the repressive function of MATR3 involves formation of a multiprotein complex with PTBP1 and possibly additional LINE-binding RBPs. Indeed, MATR3 has been reported as part of several nuclear multimeric complexes (Damianov et al., 2016, Kula et al., 2011, Zhang and Carmichael, 2001). One of these is the LASR complex, which includes hnRNPM (Damianov et al., 2016), an RBP we find preferentially binds to young antisense L1 elements much like MATR3. Taken together, our results suggest that L1 elements are sites of multivalent binding of PTBP1 and possibly other RBPs. This can provide the high avidity for assembly of repressive ribonucleoprotein complexes in order to insulate the antisense L1s and nearby RNA from RNA processing machineries.

### LINE-derived exons are highly tissue-specific

Evolutionary young LINE insertions are bound by a large number of splice repressors. This is the likely reason why only small numbers of LINEs form canonical exons, even though most LINEs contain strong 3’ and 5’ splice site sequences. Based on the often ubiquitous expression of MATR3, hnRNPM and other repressive RBPs across human tissues (Petryszak et al., 2016), LINEs need to escape this repression before they can be spliced into exons. In agreement with this, we found that LINE-derived exons are alternatively spliced, with large differences in inclusion between tissues and often completely absent from most of them. For instance, 398 of the 1,169 detectable LINE-derived exons are restricted to less than five tissues in humans; and in wildtype adult mouse brain, where activity of PTBP1 and PTBP2 is decreased, only a handful of LINE-derived exons become included at 10% or higher. However, we found strong de-repression of LINE-derived exons in PTBP2^−/-^ mouse brain and in cultured human cells in the absence of MATR3 and PTBP1. Since many different RBPs bind to intronic LINEs, it is likely that the regulation of LINE elements is combinatorial, such that the abundance of multiple RBPs determines inclusion of LINE-derived exons in each tissue. In addition, most PTBP1-repressed LINE-derived exons trigger NMD, which is likely to be common across evolutionarily young LINE-derived exons. Hence, a *bona fide* LINE-derived exon has to overcome both the splicing repressive mechanisms and NMD in order to form alternative, tissue-specific exons.

### LINEs facilitate evolution of RNA processing

To understand the forces that drive the evolutionary emergence of new LINE-derived alternative exons and poly(A) sites, we asked how the evolutionary age of LINEs affects their distribution in vicinity to exons, and the binding patterns of RBPs as surveyed by iCLIP/eCLIP. We find that intronic LINEs that recruit strong splicing repressors such as MATR3 can have repressive effects on nearby exons, which agrees with the lower inclusion of exons that are located nearby young LINEs. Conversely, young LINEs are depleted from the vicinity of constitutive exons, most likely as a result of purifying selection against the insertion of novel LINEs near existing exons, or selection of exons outside the repressive environment created by LINEs. While LINEs were known to be depleted in immediate vicinity of splice sites (Zhang et al., 2011), we now find that the extent of this depletion is distinctly dependent on their evolutionary age.

Our analysis of RBP binding patterns on LINEs demonstrates the difference in RNP assembly at evolutionarily young and old LINEs, which mediates their functional differences. The binding of repressive RBPs is most enriched in young LINEs, whereas binding of known splicing enhancers belonging to U2AF, TIA and SR families, and CPSF machinery (CSTF2 and CPSF6), is either increased or unchanged in the older LINEs. Hence, repressive RBPs prevent recognition of cryptic splice sites in thousands of young LINEs. The further the sequence of a LINE diverges, the more likely binding of one or more repressive RBPs is lost, thus allowing individual elements to seed lineage-specific and highly tissue-specific exons with low or modest inclusion. These exons become susceptible to evolutionary selection, which sets the scene for the emergence of a few *bona fide* exons with higher inclusion, seeded by evolutionarily old LINEs. The relationship between splicing repressors and LINEs is in many ways similar to the evolutionary dynamic of KAP1/KRAB transcription factors, which repress transcription at retrotransposons and confer robustness to transcriptional networks while facilitating evolutionary novelty (Thomas and Schneider, 2011, Imbeault et al., 2017).

## Conclusion

We propose that MATR3, PTBP1 and other repressive RBPs insulate the intronic LINEs to allow accumulation of cryptic elements. This could explain why strong RNA processing sites are prevalent in LINEs, and why LINEs remain cryptic without any deleterious effects. Evolutionarily young LINEs form a main hub for the recruitment of repressive RBPs, which in turn demarcate introns by insulating cryptic elements both within and around the LINE from processing machineries. These repressive RBPs are crucial to protect gene expression from the cryptic exons derived from LINE insertions, and it appears that a network of more than two dozen of LINE-binding RBPs contribute to this repression, and some of them possibly in a cooperative manner. We note this aligns with (1) previously proposed models of exon emergence (Modrek and Lee, 2003, Xing and Lee, 2006, Attig et al., 2016), in which lowly included alternative exons are the test bed for emergence of new exons; as well as (2) the proposal that genomes accumulate cryptic variation between lineages which is only apparent upon perturbation (Ward et al., 2013, Payne and Wagner, 2015, Tirosh et al., 2010). The consequences LINEs have on the transcriptome are apparent in the evolution of novel transcript isoforms, and the frequency of hereditary diseases occurring if one of these elements is unmasked.

## METHODS

### Cell culture and siRNA transfection

Hela and HEK293T cells were maintained in DMEM with 10% FBS at 37°C with 5% CO2 injection, and routinely passaged twice a week. Cells were regularly cultured for three days in antibiotic-free medium and tested for mycoplasma using either the LookOut Mycoplasma PCR Detection Kit or the MycoAlert mycoplasma detection kit (Lonza).

To deliver siRNAs, Lipofectamin RNAiMax (Life Technologies) was used according to manufacturer’s recommendations. HeLa cells were grown in antibiotic-free medium and forward transfected with siRNA targeting PTBP1 at 10nM (AACUUCCAUCAUUCCAGAGAA) and PTBP2 at 5nM (AAGAGAGGAUCUGACGAACUA), synthesized by Dharmacon, or siRNAs purchased from Invitrogen targeting MATR3 mRNA at 5nM (HSS114732) or 20nM (HSS114730), as well as control siRNA Negative Control with medium GC content (20nM, Invitrogen, Cat. number 12935-300).

### Nucleo-cytoplasmic fractionation

Cells were washed ice-cold PBS and lysed with NP40E-CSK (350μl per well of a 6-well-plate or 600μl per 10cm dish). NP40E-CSK buffer is similar to the cytoskeleton buffer used in (Reyes et al., 1997) and composed of 50 mM Tris-HCl (pH 6.5), 100 mM NaCl, 300 mM sucrose, 3mM MgCl2, 0.15% NP40 and 40 mM EDTA). Lysis was allowed to proceed for 5 minutes on ice. HeLa cells had to be scraped off due to their strong adhesion to the culture dish. Cytoplasmic supernatant and pelleted nuclei were separated at 4°C, 5000 × g for 3 minutes. The cytoplasmic supernatant was cleared with another spin at 4°C, 5000 × g for 3 minutes and another spin at 4°C, 10000 × g for 10 minutes. Nuclei were washed with 400μl NP40E-CSK and incubated for 5 minutes under rotation to ensure complete cell lysis. After repeat of the centrifugation step, nuclei were lysed in 300μl CLIP lysis buffer and sonicated at 5x 30sec pulses in a BioRuptor waterbath device. Subsequently, RNA was isolated using Trizol LS (Invitrogen) and Zymo RNA isolation columns (Zymogen) according to manufacturer’s recommendations. For preparation of RNA for RNAseq, an additional wash step with 180μl NP40E-CSK was done before nuclei rupture.

### Semi-quantitative RT-PCRs

Reverse transcription was done with 500ng of RNA using RevertAid enzyme (Fermentas) according to manufacturer’s recommendations. The reverse transcription was primed with equal parts of random N6 and N15 oligonucleotides (Sigma) at 100μM concentration. For semi-quantitative PCR, we run 35 cycles of amplification with the primer combinations as indicated in each figure (primers are listed in Supplementary Table 1), and quantified the abundance of each product using Qiaxcel™ (Qiagen) gel electrophoresis.

### UV crosslinking assay on recombinant proteins

For in vitro assays, we made two artificial sequences. The first contained two embedded AUCUU motifs (shown in bold) and CT-rich stretches in their vicinity (underlined):

GAATACGAATTCCATATATGATCGATAAATATATGGTA<u>CCTT</u>G<u>CT</u>**ATCTT**AC**ATCTT**<u>TTT</u>ACGGA<u>TCCC</u>ATATATGATCGATATATATAAGCT.

The second RNA probe contained six CTCC motifs (shown in bold):

GAATACGAATTCC**CTCTT**TGAATCGATAA**CTCTT**TGGTACCC**CTCTT**TGATCGATAA**CTCTTTGGATCCC**CTCTT**TGATCGATCTCTT**TAAGCTT.

The RNA probes were labelled with ^32^P-UTP using SP6 RNA polymerase. We purified full-length N-terminal His-tagged recombinant PTBP1 (rPTBP1) and three MATR3 fragments (rMATR3, amino acids 362-592 or ‘RRMs’, and amino acids 341-592 or ‘RRM-PRI’ with or without mutations in the PRI motif), using Blue Sepharose 6 and HisTrap HP columns. In UV crosslinking assays with recombinant proteins, we used 10fmol of RNA, 0.5μM rPTBP and titrated increasing amounts of rMATR3 fragments against it (0 to 2 μM). After incubation at 30°C for 20 minutes, the sample was UV cross-linked on ice in a Stratalinker with 1920 milliJoule. The binding reaction was then incubated for 10 minutes at 37°C together with 0.28 mg/ml RNase A1 and 0.8 U/ml RNase T1. SDS loading buffer was added and the samples heated to 90°C for 5 minutes before loading on 15% denaturing polyacrylamide gel. To assay binding in HeLa nuclear extract, we prepared standard nuclear extract (Dignam et al., 1983), and combined 10fmol of RNA probe with 0.5 μM rMATR3 and 20% extract.

### Generation of iCLIP data

HEK293T cells were grown on 10 cm^2^ dishes, incubated for 8 h with 100 μM 4SU and crosslinked with 2x 400mJ/cm^2^ 365nm UV light. Protein A Dynabeads were used for immunoprecipitations (IP). 80 μl of beads were washed in iCLIP lysis buffer (50 mM Tris-HCl pH 7.4, 100 mM NaCl, 1% NP-40, 0.1% SDS, 0.5% sodium deoxycholate). For the preparation of the cell lysate, 2 million cells were lysed in 1 ml of iCLIP lysis buffer (50 mM Tris-HCl pH 7.4, 100 mM NaCl, 1% NP-40, 0.1% SDS, 0.5% sodium deoxycholate) buffer, and the remaining cell pellet was dissolved in 50 μL MSB lysis buffer (50mM Tris-HCl pH 7.4, 100mM NaH2PO4, 7M UREA, 1mM DTT, (Reyes et al., 1997)). After the pellet had dissolved, the mixture was diluted with CLIP lysis buffer to 1000 μl and an additional centrifugation was performed. We found by Western Blotting that up to 50% of MATR3 protein is insoluble by detergent without urea. Lysates were pooled (2ml total volume) and incubated with 4 U/ml of RNase I and 2 μl antiRNase (1/1000, AM2690, Thermo Fisher) at 37°C for 3 min, and centrifuged. We took care to prepare the initial dilution of RNase in water, since we found that RNase I gradually loses its activity when diluted in the lysis buffer. 1.5 ml of the supernatant was then added to the beads and incubated at 4°C for 4 h. The rest of the iCLIP protocol was identical to the published protocol (Huppertz et al., 2014).

### Mapping of iCLIP and eCLIP data

MATR3 and PTBP1 iCLIP libraries were sequenced on Illumina HiSeq2 machines in a single-end manner with a read length of 50 nt. Before mapping the reads, we removed adapter sequences using the FASTX toolkit version 0.7 and we discarded reads shorter than 24 nucleotides. Reads were then mapped with the iCount suite to UCSC hg19/GRCh37 or mm9/NCBI37 genome assembly using Bowtie v2.0.5 allowing up to two mismatches and up to 20 multiple hits. Unique and multiple mappers were separately analysed, and to quantify binding to individual loci, only uniquely mapping reads were used. Supplementary Table 1 lists the source and details including fil numbers of all published iCLIP and HITS-CLIP data used within this study.

The eCLIP libraries were downloaded from ENCODE (Van Nostrand et al., 2017, Sloan et al., 2016). Before mapping the reads, adapter sequences were removed using Cutadapt v1.9.dev1 and reads shorter than 18 nucleotides were dropped from the analysis. Reads were mapped with STAR v2.4.0i (Dobin et al., 2013) to UCSC hg19/GRCh37 genome assembly. To quantify binding to individual loci, only uniquely mapping reads were used. For analysis of enrichment on repeat families, up to 20 multiple alignments were allowed and fractional counts used.

### TEtranscript estimates of LINE family enrichments

To consider both uniquely mapping and multimapping reads in estimating binding to repeat (sub)families, we used the approach described in TEtranscripts (Jin et al., 2015). In short, for eCLIP FASTQ files, adapters were removed according to the ENCODE eCLIP standard operating procedure. For iCLIP FASTQ files, barcodes were removed using the FASTX-Toolkit (v 0.0.14). For all files, reads aligning to rRNA or tRNA were removed by aligning to custom rRNA and tRNA indices (human or mouse as appropriate) using Bowtie2 (v. 2.2.9). The remaining reads were aligned to the appropriate genome (GRCh38, Gencode V25 for human, and GRCm38, Gencode M13 for mouse) using STAR (v. 2.5.2) with the addition of the parameters “`-- winAnchorMultimapNmax 100 --outFilterMultimapNmax 100`” as recommended by TEtranscripts. For each CLIP dataset, TEtranscripts was run using both stranded options (`--stranded reverse` and `--stranded yes`) to obtain results for sense and antisense LINE binding.

RNAseq data from ENCODE was used as control, for eCLIP RNAseq of K562 and HEPG2 cells lines (ENCSR885DVH and ENCSR181ZG). For iCLIP samples from mouse brain, we used P2 mouse brain from ENCODE. The iCLIP data in mouse brain was produced from total mouse brain, so we pooled the RNAseq of forebrain, midbrain and hindbrain, accession numbers ENCSR723SZV, ENCSR255SDF and ENCSR749BAG (Sloan et al., 2016).

### Generation of RNAseq libraries and mapping with TopHat2 (human)

Before library preparation, purified RNA was DNase I treated for a second time and purified with the DNA-free kit (Ambion). To generate stranded RNAseq libraries, we used the TruSeq stranded RNAseq library kit (Illumina) according to manufacturer’s recommendations; RNA was depleted of rRNA using the RiboZero kit (Epicentre).

All libraries were sequenced on Illumina HiSeq2 machines in a single-end manner with a read length of 100 nt. Before mapping the reads, adapter sequences were removed using the FASTX toolkit version 0.7 and we discarded reads shorter than 24 nucleotides. Reads were then mapped with TopHat v2.0.5 (Kim et al., 2013) to UCSC hg19/GRCh37 genome assembly using ENSEMBL version 72 gene annotation as reference, allowing up to two mismatches and only using uniquely mapping hits. RNAseq data files of rRNA depleted cytoplasmic and nuclear RNA from cells depleted of MATR3 and PTBP1 are deposited on EBI ArrayExpress under the accession number E-MTAB-6204.

### Generation of pAseq libraries and mapping

To quantify polyA site usage, we used the QuantSeq mRNA 3’ end sequencing kit (Lexogen) according to manufacturer’s recommendations. We used both the forward and reverse library kit on two independent biological replicates each (four replicates in total). Libraries were prepared from nuclear RNA after individual or combined siRNA depletion of MATR3 and PTBP1/2. All libraries were sequenced on Illumina HiSeq2 machines in a single-end manner with a read length of 100 nt. polyA site usage was analysed with the expressRNA platform. In short, reads were mapped with STAR v??? to UCSC hg19/GRCh37 genome assembly, allowing up to ??? mismatches and ?only using uniquely mapping hits?. pAseq raw data is deposited on ArrayExpress at E-MTAB-6287.

### Mapping of published RNAseq with STAR (human)

To test for any change in usage of LINE-derived exons upon depletion of the NMD core factor UPF1, we made use of the data generated by Ge et al. and in HEK293 cells, depleted of PTB, UPF1 or both proteins (Ge et al., 2016). Raw sequencing data in FASTQ format was downloaded from SRA and mapped with STAR v2.5.2a (Dobin et al., 2013) to UCSC hg19/GRCh37 genome assembly, allowing up to 10 mismatches and only using uniquely mapping hits. Then we analysed the data using JunctionSeq (Hartley and Mullikin, 2016) with ENSEMBL version 72 gene annotation as reference.

### Mapping of published RNAseq with STAR (mouse)

To test for LINE-derived exon inclusion in mouse brain, we made use of the data generated by Li et al. and Vuong et al. (Li et al., 2014, Vuong et al., 2016). Raw sequencing data in FASTQ format was downloaded from SRA and mapped with STAR v2.5.2a (Dobin et al., 2013) to UCSC mm10/GRCm38 genome assembly, allowing up to 10 mismatches and only using uniquely mapping hits. Then we analysed the data using JunctionSeq (Hartley and Mullikin, 2016) with ENSEMBL version 72 gene annotation as reference.

### Sequence motif analysis

For PTBP1 motifs around iCLIP peaks, we used the strong binding motifs as defined previously (Haberman et al., 2017), and counted their occurrence around peak centres. To define enrichment, we divided the occurrence at MATR3-dependent and independent peaks by the distribution across all other PTBP1 peaks.

For motifs within antisense L1 elements, we used motifs described in the literature; for PTBP1, TARDBP and hnRNPM their binding motifs were validated in vitro and through functional studies (Gooding et al., 1998, Oberstrass et al., 2005, Avendano-Vazquez et al., 2012, Ayala et al., 2005). For all other proteins, we used RNAcompete motifs (Ray et al., 2013). The number of motifs per 100nt gave a distribution for each motif, and we used quartiles for each motif to describe gain or loss of motifs between evolutionary groups. To obtain a false discovery rate of motif gain or loss, we generated an empirical distribution of motif enrichments across groups. We compared the change in Q1 and Q4 for each of the possible 256 tetramers, which resulted in an approximately normal distribution. We then called motifs within the 2.5% and 5% extremes as significant at FDR<0.05 and FDR<0.1.

### RNA maps

All metaprofiles of iCLIP data and LINE sequence content around loci of interest (also called RNAmaps) were drawn in R. Metaprofiles are normalised to the number of input loci of each track, and data was smoothed using binning as indicated in figure legends, using the *zoo* package. A generalised script for generation of a metaprofile can be found at https://github.com/JAttig/generalised-Rscripts.

### Annotation of known alternative exons

For annotation of exons known to be alternatively spliced, we downloaded the ‘knownAlt Events’ and ‘knownGene’ from UCSC TableBrowser for hg19 on 28^th^ March 2014. In addition, we downloaded the ‘refGene’ table on 23^rd^ March 2017. The exons annotated by UCSC were collapsed within a gene to unique exonic ranges, and classified as alternative or constitutive exon as follows. All exons not annotated as alternative by UCSC and present in the RefSeq exon annotation with identical genomic coordinates were classified as constitutive, all other exons were considered alternative exons.

### *De novo* identification of cryptic exons

In order to predict exons from our RNAseq data, we ran Cufflinks (version 0.9.3,-min-isoform-fraction 0, Trapnell et al., 2012) on the collapsed reads from all cytoplasmic samples of our stranded RNAseq data and then extracted the exons of all predicted transcripts. After flattening the Cufflinks output to non-overlapping exonic bins, our Cufflinks prediction contained 671,956 exonic bins. However, we only considered exonic bins of at least 5 nucleotides in size. All exons with one or both splice sites residing within a LINE repeat (as annotated by RepeatMasker, (Smit et al., 1996-2010b)) were assigned as LINE-exons. In order to minimise noise, we only kept exons for analysis that were supported by at least one junction-spanning read (225,322 exonic bins). All exons that were not identical with exons annotated in UCSC gene annotation (hg19) were referred to as ‘cryptic’ (see also Supplementary Table 2) for complete breakdown of annotation of exonic bins). For readability, we refer to ‘exonic bins’ as ‘exons’ throughout the text.

### Analysis of differential gene expression and differential exon inclusion

Analyses of differential gene expression were performed using DESeq2 (Anders and Huber, 2010, Love et al., 2014) with gene coordinates based on ENSEMBL annotation (version 72). To combine the results from both siRNAs targeting MATR3, we used a conditional thresholding approach, calling expression changes as significant if they had an adjusted p-value < 0.01 in at least one of the two depletion conditions and an adjusted p-value < 0.05 in the other. Differential splicing was determined using DEXSeq (Anders et al., 2012), and the two MATR3 depletion conditions were combined by conditional thresholding accordingly.

### Analysis of exon inclusion in human tissues

To analyse inclusion of exons across human tissues, we retrieved data on mapped junctions from the V6p release of the GTEx consortium (http://www.gtexportal.org/home/, (Consortium, 2015)). We used UCSC/RefSeq annotation (see above) and isolated all LINE-derived exons as well as Alu-exons. Then, we selected all exons from genes with at least one Alu- or LINE-derived exon. We identified junction-spanning reads to each of these exons in a 2 nt grace window around the splice and used those to identify the 5’ and 3’ splice site of the upstream and downstream exon. We then calculated percent spliced in (PSI) index as the ratio of inclusion junction reads (average of up+downstream junctions) to total junction reads (average of up+downstream junctions + skipping junctions), and inclusion within each tissue as average of all samples. We only allowed a single exon inclusion isoform across tissues (i.e. identical flanking exons) and choose the isoform with more junction reads. To ensure sequencing depth and gene expression were sufficient to calculate exon inclusion, we only used exons with at least 200 reads across the 8,555 samples (average of up+downstream junctions or skipping junctions). If an exon was absent in any tissue, as judged by absence of any junction spanning read and any read for the skipping junction, it was treated as ‘data not available’ for this particular tissue. In total, we covered 45,940 exons across 52 tissues and subtissues, which were adipose tissue (subcutanoues and visceral omentum), adrenal glands, artery (aorta, tibial and coronary artery), bladder, brain, breast, cervix (ecto- and endo-cervix), colon (sigmoid and transverse), esophagus (mucosa, muscularis and gastroesophageal junction), fallopian tube, heart (atrial appendage and left ventricle), kidney (cortex), liver, lung, skeletal muscle, nerve tissue (amygdala, anterior cingulate cortex, caudate basal ganglia, cerebellar.hemisphere, cerebellum, cortex, frontal cortex, hippocampus, hypothalamus, nucleus accumbens basal ganglia, putamen basal ganglia, cervical spinal cord, substantia nigra, tibial), ovary, pancreas, pituitary, prostate, minor salivary gland, small intestine terminal ileum), spleen, skin (suprapubic and lower leg), stomach, thyroid, testis, uterus and vagina, as well as EBV transformed lymphocytes and transformed fibroblasts. We did not use data from whole blood, which had poor coverage on most genes. On top of the PSI index for each tissue, we collated the data across tissues and computed the maximum difference in PSI between the tissue(s) with highest inclusion and lowest inclusion of each exon. Because testis is known to be a very promiscuously transcribed tissue (Soumillon et al., 2013) and accordingly showed many LINE-derived exons exclusively observed in the testis, we only included exons which showed at least 5% inclusion in any tissue, except testis.

### Classification of repeat element age by divergence or phylogenetic tracing

To compare the divergence of LINE insertions from their consensus sequence, we used the nucleotide difference / 1000nt, which is provided for each repeat element by the RepeatMasker table (hg19, Repeat Library 20090604, (Smit et al., 1996-2010a)).

For phylogenetic tracing, we tested for presence of orthologues positions with the UCSC Genome Browser LiftOver tool (Rosenbloom et al., 2015), using the respective all-chain BLASTZ files. Human and mouse LINE repeats from hg19 and mm9 RepeatMasker annotation were first lifted to hg38 and mm10. We then tested for the presence of each LINE repeat in the human and mouse lineage by retrieving orthologue genomic loci for the genomes of rhesus macaque (rheMac8), gorilla (gorGor5), mouse (mm10), rat (rn6), dog (canFam3) and cow (bosTau8). To curate the LiftOver results and safeguard against misannotation by errors in the genome lift, we cross-referenced for all liftover positions if the element overlaps with a LINE annotated by RepeatMasker for the respective genome, and only refer to the element as present in a species if at least 33% of the lifted genomic position are LINE-derived as annotated by RepeatMasker. All other elements are either ‘notLINE’ if they were not identified by RepeatMasker, ‘degenerate’ if LiftOver reported them as ‘partially-deleted’, or ‘absent’ if LiftOver reported them as ‘deleted’. Elements from hg19 that were not ‘present’ in hg38 were discarded entirely. Then we converted the LiftOver annotation to phylogenetic groups after manual inspection of the liftover results in the following manner. We denoted elements as human- and primate-specific, which are ‘absent’ in all other species. We denoted additional elements as primate-specific, if they were either ‘present’, ‘degenerate’ or ‘notLINE’ in at least one of the two primate species, and ‘absent’ or ‘notLINE’ in all of the others. We denoted elements as specific for the euarchontoglires branch, if the element was ‘absent’ or ‘notLINE’ in the two laurasiatherian species, and ‘present’ or ‘degenerate’ in mouse or rat. The remaining elements were all lifted towards at least one of the two laurasiatherian species, and hence present in the last common ancestor of the species we surveyed. Elements present in one but absent in the other were denoted as found in ‘one distant species’, elements present in both as found in ‘two distant species’. All remaining elements were either reported as degenerate in both species, or the liftover results were ‘unclear’ (for example if the element was lifted to many species but did not overlap with the LINE annotation in any of those). In either case, we ignored the corresponding element for phylogenetic comparisons. Group sizes for the hg19 assembly were:

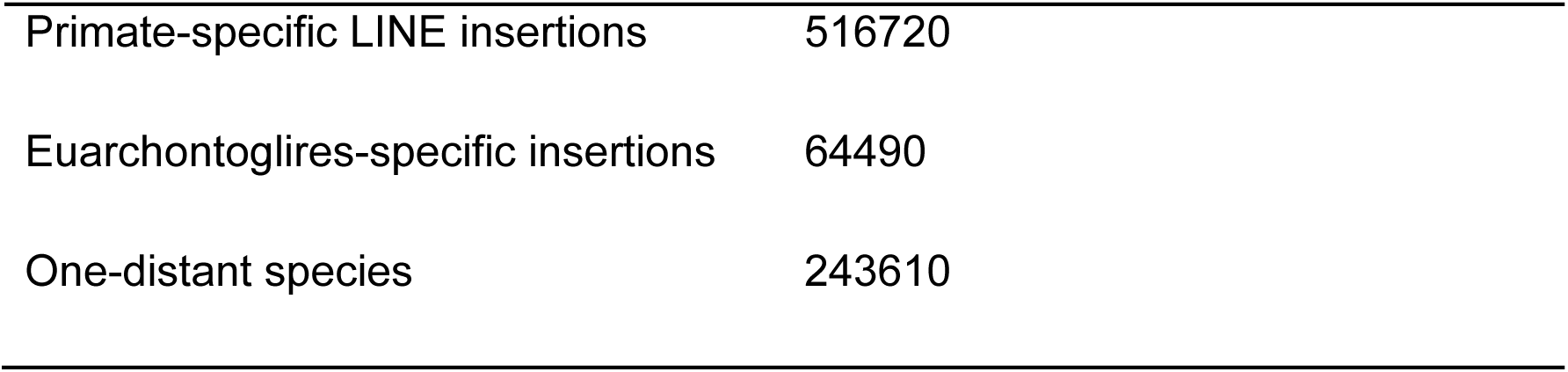

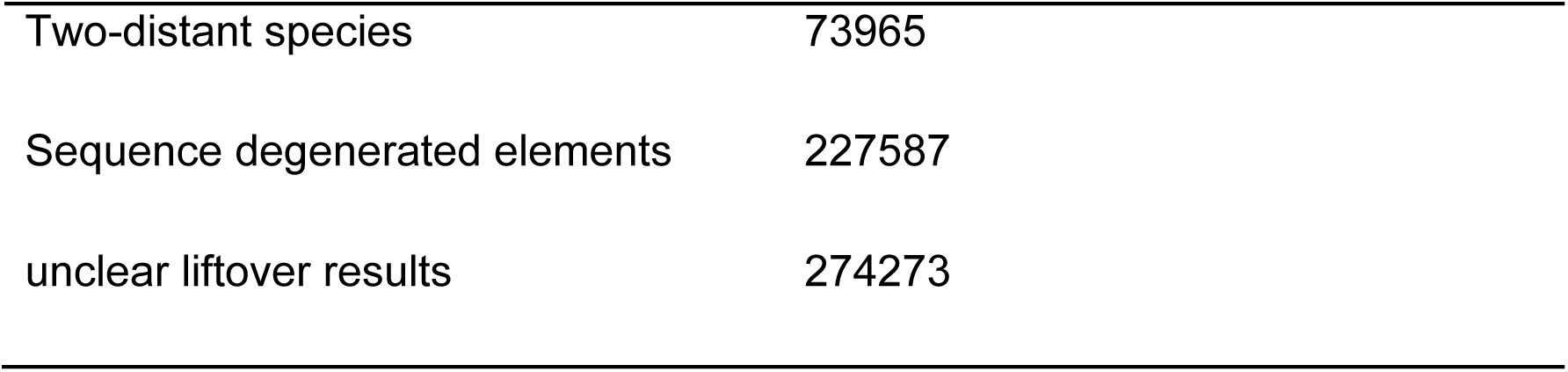

## Statistics

Whenever referred to in the text, *replicates* stands for biological replicates, defined as samples collected independently of one another in separated experiments. All experiments were done with biological replicates as indicated in Methods and Figure legends. In case of the iCLIP experiments from MATR3 or PTBP1 depleted cells, sequencing files were pooled across 2 biological replicates because coverage varied widely within them, and only the pooled data was used.

All statistical analyses were performed in the R software environment (version 3.1.3) or in GraphPad PRISM6. We made use of nonparametric tests in all statistical tests, since data distributions failed to conform with the assumption of normality and equal variance (homoscedasticity), assessed visually with qqnorm plots. Statistical tests are listed in figure legends. To compare multiple groups we used the Kruskal-Wallis Rank Sum test, and made pairwise comparisons with Dunn’s test corrected according to Holm-Sidak, using functions implemented in the *stats* and the *dunn.test* (Dinno, 2017) R packages.

## ACKNOWLEDGEMENTS

The authors are grateful to Michael Briese, Laura Easton, Ina Huppertz and James Tollervey for sharing unpublished iCLIP data. We thank Ina Huppertz and Igor Ruiz de los Mozos for assistance and scientific discussion, Sarah K. Jurmeister for critical comments on this manuscript and Gavin Kelly for valuable advice. We thank the Genomics Facility Team of the CRUK Cambridge Cancer Institute and Deborah Hughes of the UCL Institute of Neurology for processing libraries for high-throughput sequencing, and Gregor Rot for mRNA 3’ end sequencing mapping on the expressRNA platform. This work was supported by the European Research Council (617837-Translate to J.U.), the Wellcome Trust with a Wellcome Trust Joint Investigator Award (103760/Z/14/Z; to J.U. and N.L.), a Wellcome Trust Programme grant (092900; to CWJS) and a PhD Training Fellowship for Clinicians Award (110292/Z/15/Z; to A.C.), and a Boehringer Ingelheim Fond PhD fellowship (to J.A.). This work was supported by the Francis Crick Institute, which receives its core funding from Cancer Research UK (FC001110), the UK Medical Research Council (FC001110), and the Wellcome Trust (FC001110) (N.L., J.A., F.A., A.C.). Animal shapes in Figure 5 were obtained from PhyloPic and are used under the Creative Commons Attribution-NonCommercial-ShareAlike 3.0 Unported license. Images created by Michael Kessey, David Liao and Maija Karala.

## AUTHOR CONTRIBUTIONS

J.A., C.G., C.W.J.S and J.U. conceived the project and designed the experiments. F. A. supervised computational analysis. J.A., C.G. and A.S. performed experiments, and J.A., F.A., A.C., N.H. and W.E. performed computational analysis. J.A., F.A, C.S., N.L. and J.U. interpreted and conceptualised primary data.

## List of Supplementary files and Tables

**Suppl. Table 5: Annotation derived from phylogenetic tracing of LINE elements in hg19**.

**Suppl. Table 6: Inclusion levels of 43583 UCSC annotated exons in 53 human tissue types**.

**Suppl. Table 7: Summary statistics of tetramer frequencies in antisense L1 sequences**.

## FIGURES AND FIGURE LEGENDS

**Figure S1.**
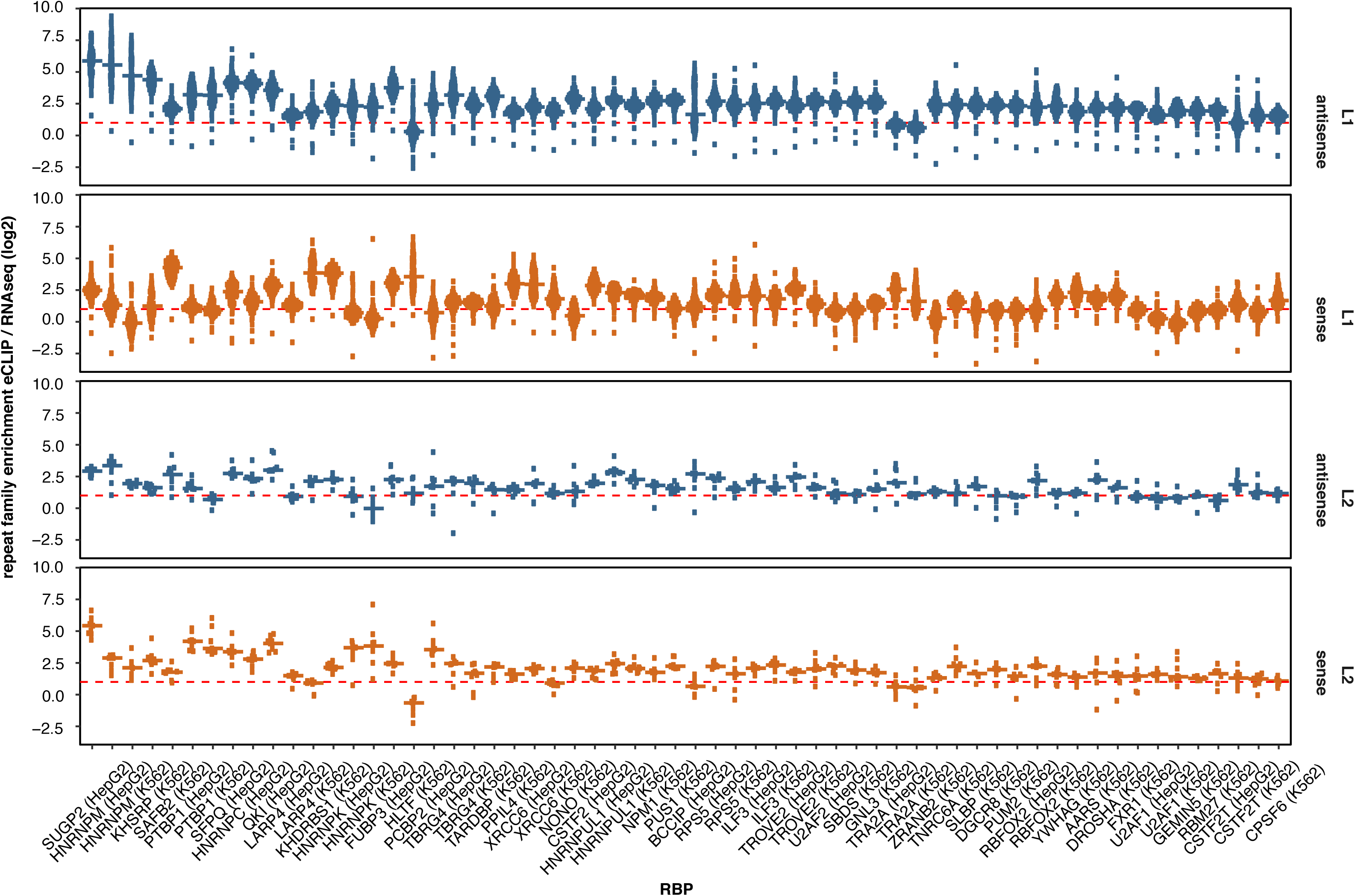
Related to Figure 1: Extended data for LINEs are binding platforms for a set of RBPs. (A) TEtranscript (Jin et al., 2015) was used to estimate the enrichment of each subfamily of L1 and L2 repeats among the bound RNA sequences of a panel of RBPs, comparing the abundance in recovered eCLIP tags to the abundance in RNAseq reads. For each RBP, all 142 L1/L2 subfamilies (132 for L1, 10 for L2) were considered. Since eCLIP is strand-specific, binding to LINEs transcribed in sense or in antisense were quantified separately, coloured in red and blue. The cell lines used in each eCLIP experiment are indicated on the bottom.

**Figure S2.**
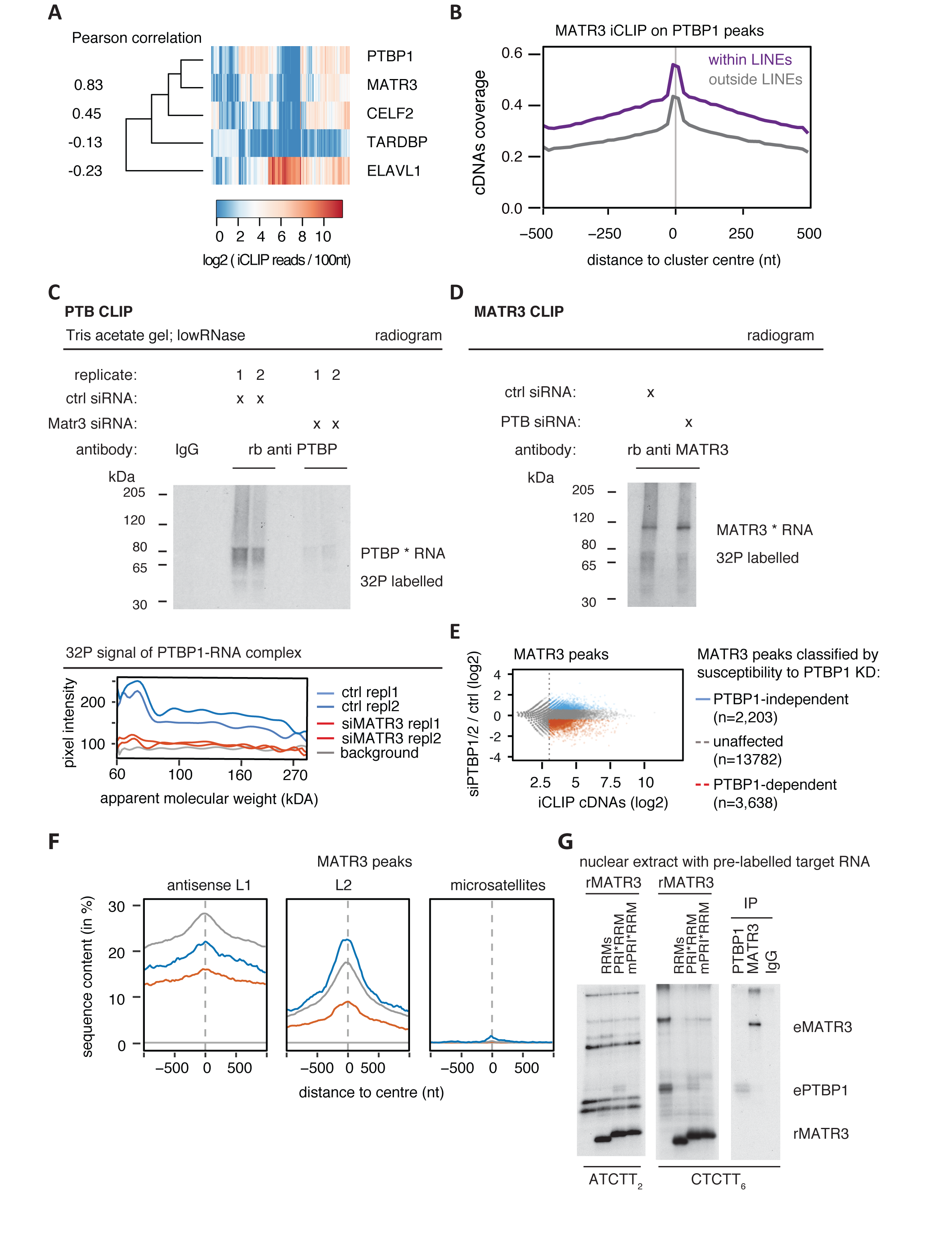
Related to Figure 2: Combinatorial binding of MATR3 and PTBP1 to the same LINEs. (A) For each RBP that showed considerable binding to LINE repeats in iCLIP (see B), we selected the 50 LINE repeats with strongest coverage (cDNAs per 100nt). For comparison we included TDP43, which showed little binding to LINE repeats. All iCLIP data selected was collected from HEK293 cells. The heatmap shows comparison of binding strength at this set of 214 LINE repeats, and the nearest neighbour analysis for each RBP. The values left to the dendrogram show the pearson correlation coefficient between all RBPs and PTBP1. Only LINEs with a minimal length of 50nt were considered to reduce the bias to short, highly expressed LINE repeats. (B) Metaprofile of iCLIP binding for MATR3 around iCLIP binding peaks of Celf2, Celf4, TDP43, HuR and PTBP1 within and outside of LINE repeats. The data was smoothed with 20nt bins. (C) HEK293T cells were transfected with siRNAs targeting MATR3, PTBP1 or scrambled controls, and 72 hours later labelled with 100μM 4SU for 8 hours and cross-linked with 365nm UV light. The radiogram shows ^32^P labelled RNA crosslinked to and co-precipitated with PTBP1. Before immunoprecipitation, protein concentration was measured and equalised. The PTBP1 iCLIP was done under low RNase conditions (compare with Fig. 2A for high RNase condition). Replicate 1 and 2 are independent biological replicates processed in parallel. (D) ^32^P labelled RNA crosslinked to and co-precipitated with MATR3 under equivalent conditions as in (C). The MATR3 iCLIP shown was done under high RNase conditions. (E) MATR3 binding peaks were identified from iCLIP experiments, and classified according to susceptibility to PTBP1 depletion as indicated based on moderated log2 fold change. Binding peaks with a normalised count of less than 8 were ignored, as indicated by the dotted line. (F) The overlap between the centre of MATR3 binding peaks and different repeat classes was tested for antisense L1 elements, sense L2 elements, and sense CT-/T-rich microsatellite repeats. Metaprofile shows percent of each class of clusters overlapping with each genomic element. (G) Protein-protein interaction between MATR3 and PTBP1 allows recruitment of PTBP1 to a MATR3 bound RNA *in vitro.* Recombinant MATR3 mutants (rMATR3) and 32P labelled RNA probes were added to nuclear extracts from HeLa cells and UV-crosslinked. RNA substrates contained either two MATR3 or six PTBP1 RNA compete motifs motifs (ATCTT_2_ and CTCTT_6_). Crosslinking signals corresponding to endogenous PTBP1 (ePTBP1) and MATR3 (eMATR3) were confirmed by immunoprecipitation.

**Figure S3.**
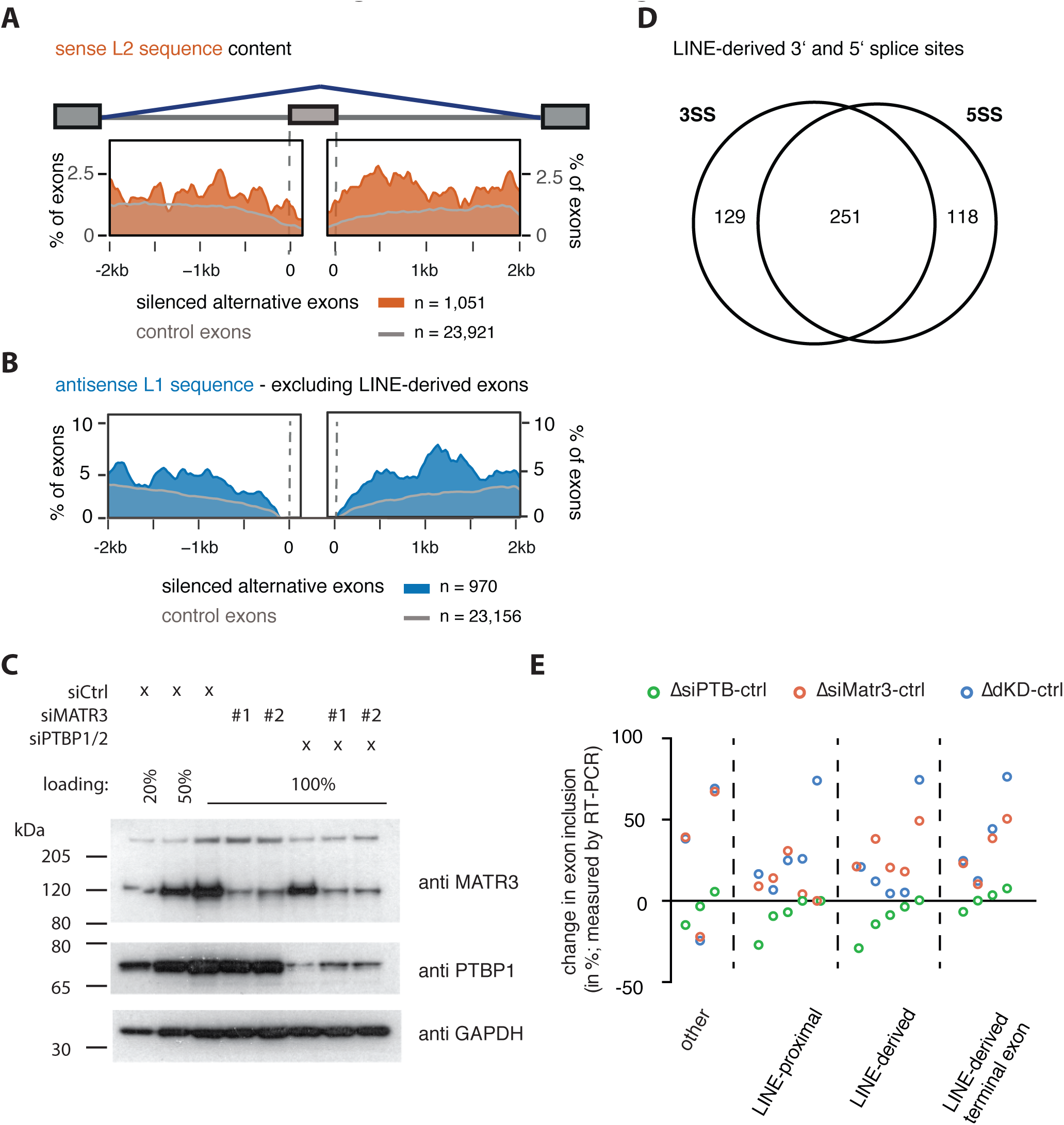
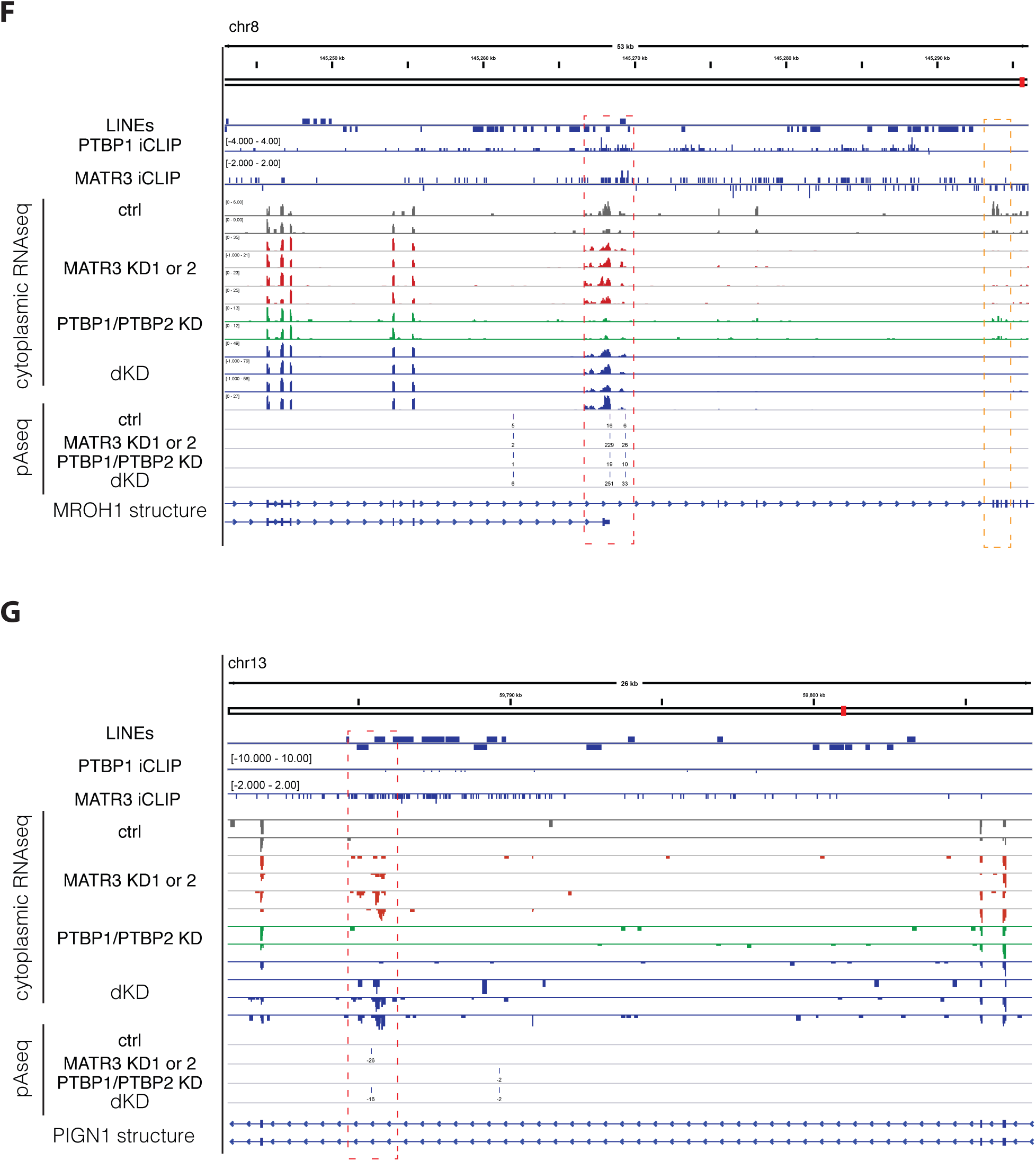
Related to Figure 3: MATR3/PTBP1 repressed exons are frequently derived from LINEs or proximal to LINEs. (A) The metaprofile shows the amount of sense L2 sequences flanking the splice sites of MATR3/PTBP1/2 repressed events. L2 sequences are particularly enriched towards the 3’ splice site, and to a lesser extent than antisense L1 sequence. (B) The metaprofile shows the amount of antisense L1 sequences flanking the splice sites of MATR3/PTBP1/2 repressed events, after removing all LINE-derived exons. The enrichment for L1 antisense sequence still persists (compare with Fig. 3A). (C) Semi-quantitative Western blot showed efficient depletion of MATR3 and PTBP1 in cells transfected with siRNAs against MATR3 or PTBP1 individually or in combination. (D) The overlap of LINE-derived exons for which the 3’ or 5’ splice site is derived from a LINE element, only showing exons with junction-spanning reads on both sides (498 exons). (E) Seventeen exons differentially included in MATR3 depleted cells were selected, and changes in exon inclusion were validated by RT-PCR. For each exon, the relative abundance of the isoform including the alternative exon was calculated compared to the exon exclusion isoform (conventional splicing pattern). The change between cells depleted of MATR3, PTBP1/2, or both simultaneously is shown, and exons are grouped by their positioning relative to the closest LINE element. Semi-quantitative RT-PCR analysis is averaged across three independent replicates. (F) And (G) Examples of MATR3/PTBP1 repressed polyA sites. Genome browser tracks show position and orientation of LINE insertion (hg19/RepeatMasker annotation), PTBP1 and MATR3 iCLIP coverage, as well as tracks for RNAseq of cytoplasmic RNA and mRNA 3’ end sequencing (pA-seq) from total RNA. All tracks are scaled appropriately to library size. (F) The MROH1 gene shows inclusion of additional exonic sequence and two different terminal exon isoforms in MATR3 depleted cells (highlighted by red dashed lines). Usage of this alternative terminal exon results in gene truncation as seen by loss of expression downstream of it (highlighted by orange dashed lines). (G) The PIGN1 shows usage of a cryptic processing site resulting in a novel exon and a novel polyA site, derived from two antisense L1 insertions (highlighted by red dashed lines).

**Figure S4.**
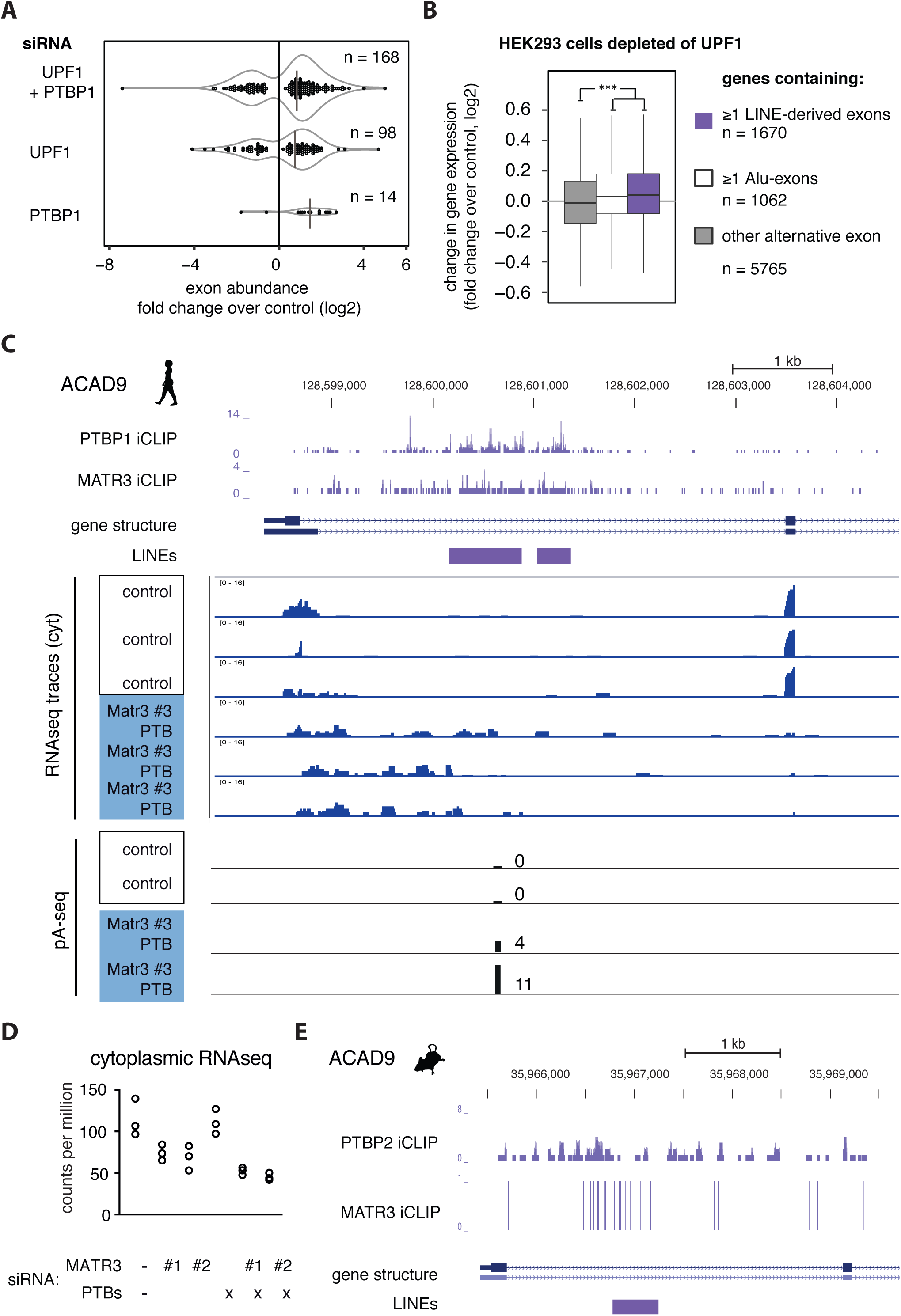
Related to Figure 4: Nonsense-mediated decay triggered by LINE-derived exons and depletion of ACAD9 expression following inclusion of a LINE-derived exons. (A) RNAseq data from Ge et al. on HEK293 cells depleted of PTBP1, UPF1 or both, was reanalysed with DEXSeq. The number of detectable LINE-derived exons and their change in abundance compared to control cells is shown. Consistent with the hypothesis that LINE-derived exons are repressed in wild-type cells by splicing repressors and through decay of the inclusion isoform, combined depletion of UPF1 and PTBP1 greatly increases the number of detectable LINE-derived exons. (B) The change in gene expression in UPF1-depleted cells over control is shown for genes that contained or did not contain one or more LINE-derived exons. As positive control, Alu-exon containing genes are shown since inclusion of Alu-exons frequently triggers NMD (Attig et al., 2016). (C) Genome browser tracks for PTBP1 and MATR3 iCLIP data from HeLa cells at the ACAD9 locus. Position of L2 insertions is annotated below the structure of annotated ACAD9 transcripts, and stranded RNAseq data from cytoplasmic RNA of HeLa cells depleted of MATR3/PTBP1/PTBP2 is shown. Below the position of a novel pA site within the second L2 repeat is shown, which is only detected in absence of MATR3/PTBP1/PTBP2. (D) Quantification of ACAD9 expression in single and combined depletion of MATR3 and PTBP1/2 from cytoplasmic RNAseq. (E) Genome browser tracks for PTBP2 and MATR3 on the mouse ACAD9 locus. In mouse, there is a single, 465bp long L2 insertion.

**Figure S5.**
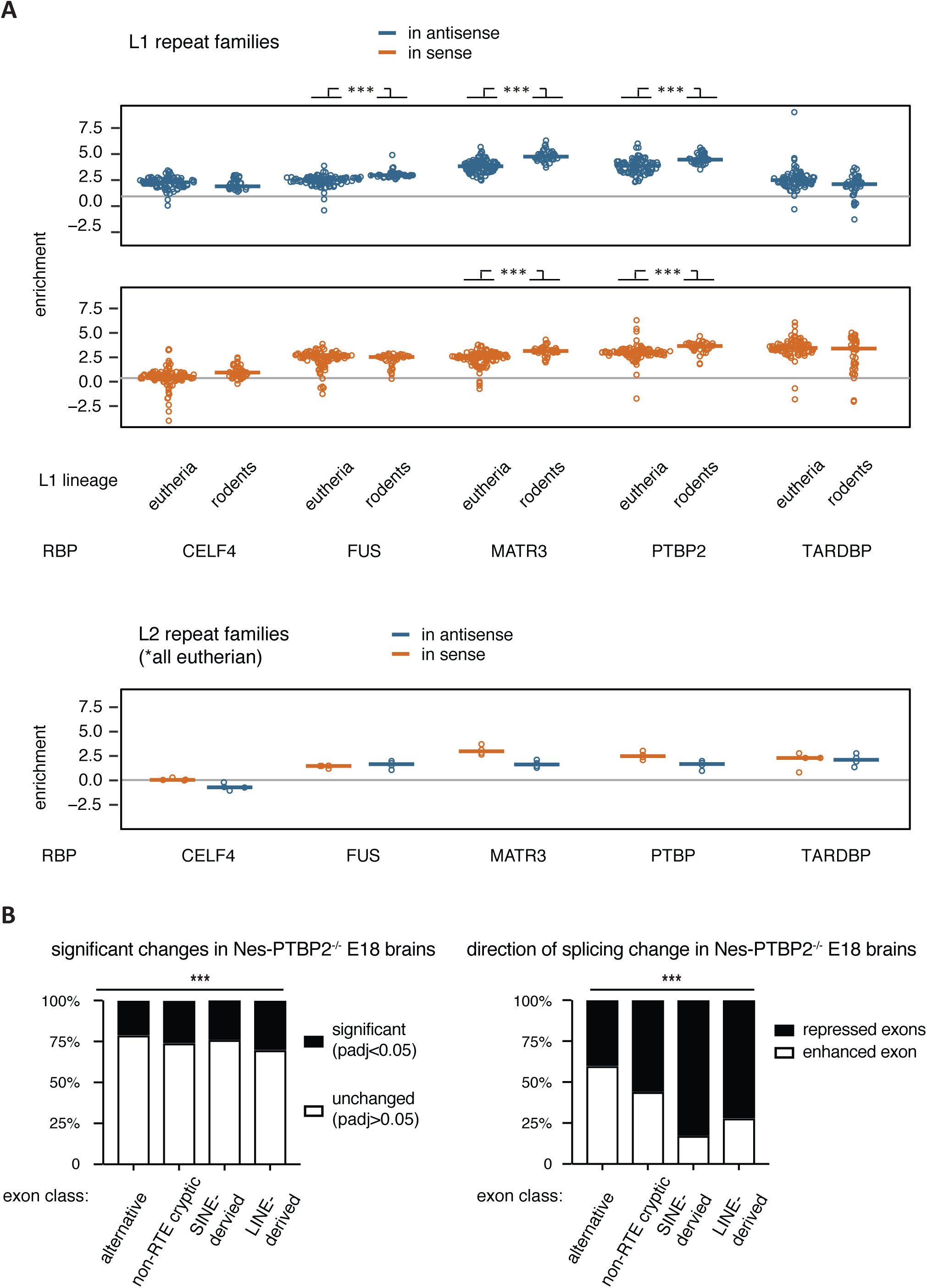
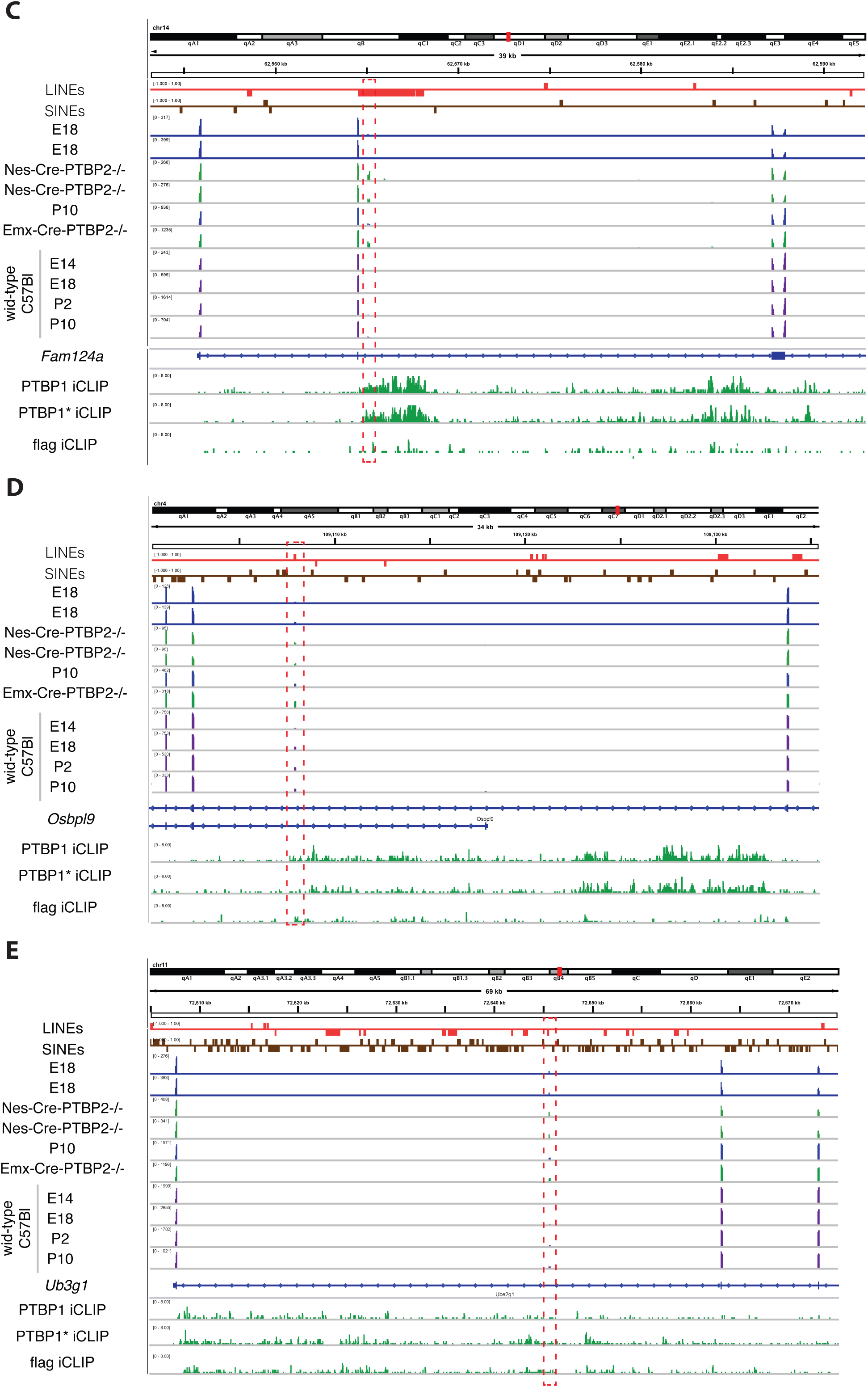
Related to Figure 5: MATR3 and PTBP2 binds to mouse-specific L1 insertions and PTBP2 represses LINE-derived exon inclusion in the mouse brain. (A) TEtranscript (Jin et al., 2015) was used to estimate the enrichment of each subfamily of L1 and L2 repeats among the bound RNA sequences of a panel of RBPs, with CLIP data available for C57Bl mouse brain; comparing the abundance in recovered eCLIP tags to the abundance in RNAseq reads of P2. For each RBP, 133 repBase LINE subfamilies were considered (129 for L1, 4 for L2, (Jurka, 1998)). Since eCLIP is strand-specific, binding to LINEs transcribed in sense or in antisense were quantified separately, coloured in red and blue. Details and references of data sets are given in Supplementary Table 1. (B) RNAseq data of PTBP2^−/-^ knockout mouse brains from (Vuong et al., 2016) was re-analysed, and exons with significant differences in inclusion in Nes-Cre-PTBP^−/-^ knockout mouse brains were stratified according to their relationship to retrotransposon repeats. LINE-derived exons were more likely to be mis-regulated than expected by chance (x^2^ test), and PTBP2 acts primarily as repressor on LINE-derived exons. Number of exons in each group are: alternative exons n=8142, non-repeat derived cryptic exons n=33420, SINE derived exons n=459, LINE-derived exons n=308. (C) to (E) RNAseq data of PTBP2^−/-^ knockout mouse brains from (Vuong et al., 2016) compared to RNAseq data of C57/B6 wild-type mouse brain at different developmental stages (E10, E14, P2 and P10). LINE-derived exons were selected from the exon list of PTBP2 responsive exons provided by Vuong et al. in Suppl Table 3 and 5; that is they are selected to show a minimum 10% change in inclusion upon PTBP2 depletion. PTBP1 iCLIP was done with flag antibody in Rosa-PTBP1 transgenic mice, and PTBP1* iCLIP is PTBP1 iCLIP in PTBP2 knockouts (also from (Vuong et al., 2016)). Because of the high extent of intron-retention reads in mouse brain, only junction-spanning reads are shown. These exons are more included in postnatal brains than in foetal brain, suggesting PTBP2 suppresses exonisation in developing neurons but less in mature neurons. (C) Exon 3 of Fam124 is derived from a rodent specific L1 insertion. (D) Exon 5 of *Osbpl9* is derived from an old CR1 insertion conserved across mammalian lineages. (E) Exon 2 of *Ube2g1* is derived from an old HAL1 insertion conserved across mammalian lineages.

**Figure S6.**
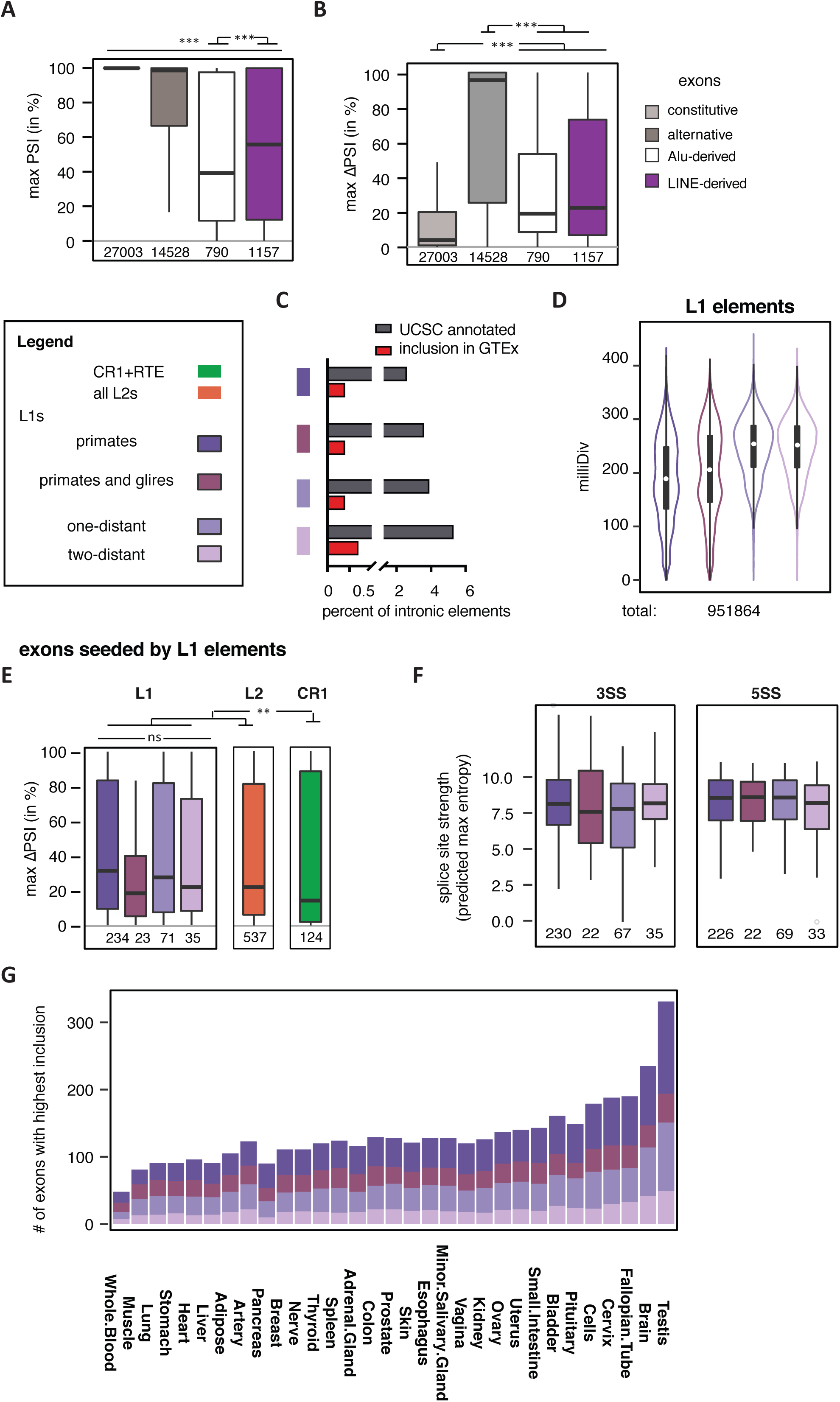
Related to Figure 5: Li-derived exons are a source of primate-specific alternative exons with high tissue-specific variability. Percent splice index (PSI) was calculated in the GTEx panel of human tissues for LINE-derived and Alu-derived exons, as well as all other exons of the same genes. All exons are annotated within UCSC and cross-referenced with RefSeq annotation. Inclusion levels range from 0 to 100%, showing no inclusion or full inclusion. If no support for expression of the flanking exons was found, the gene is assumed to be non-expressed. Genomic age of L1 elements as defined in Figure 5A. Significance tests were done across groups by Kruskal-Wallis’ test and pairwise comparisons were done with Dunn’s test and corrected according to Holm-Šidák. ** and *** indicate adjusted p-value was below 0.01 and 0.001, respectively. (A) Maximum inclusion in any tissue was calculated for each exon, and the distribution is shown for LINE-derived exons, Alu-exons as well as non-repeat derived alternative and constitutive exons. (B) For all exons surveyed within the GTEx data, the difference in PSI between the tissues with highest and lowest inclusion was calculated as metric for tissue-specific inclusion. (C) Exons derived from old L1 insertions are most likely to form an exon based on UCSC annotation. Based on GTEx data, exons derived from old L1 insertions retained in primates, cow and dog, are most likely to be included in any of the tissue types covered. (D) The substitutions from L1 consensus families is shown for L1s grouped by genomic age. As expected, young elements show fewer substitutions from consensus then old elements. (E) Difference in PSI between tissues with highest and lowest inclusion for exons derived from L1 elements grouped by genomic age of the insertion, compared to exons derived from L2 and CR1 insertions. (F) Exons derived from L1 elements have strong splice sites irrespective of the genomic age of the insertion. The maximum entropy score of 5′ and 3′ splice sites of each exon was predicted based on nucleotide sequence (Yeo and Burge, 2004). (G) The number of L1-derived exons is shown for all primary tissues screened in the GTEx data, based on testing in which tissue an exon is most included. Exons are allowed to be counted multiple times if maximum inclusion was in multiple tissues, for instance because they are constitutive.

**Table S3.**
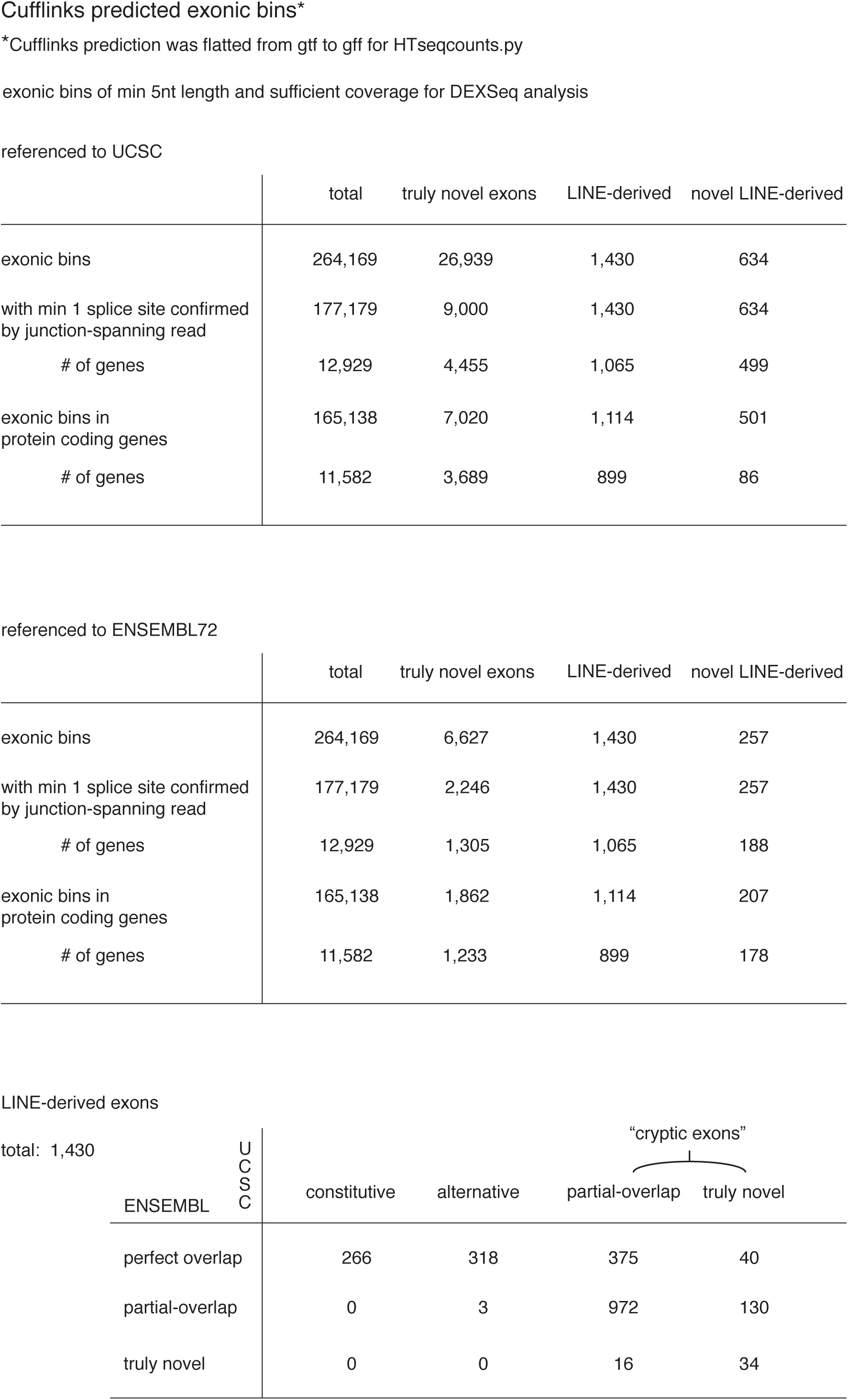
Summary statistics of cryptic exon annotation from interleaving UCSC or ENSEMBL annotation and Cufflinks assembly.

**Table S4.**
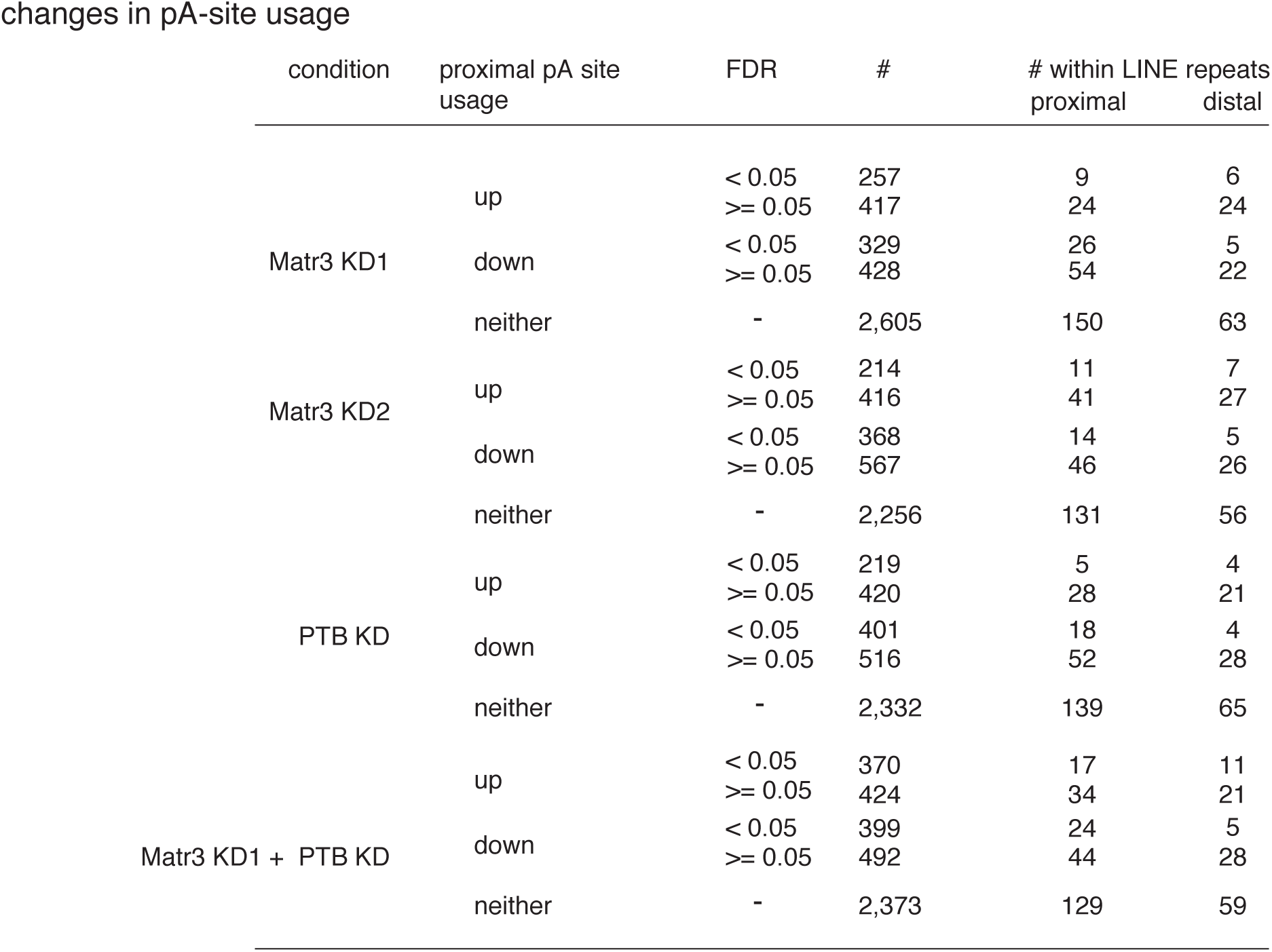
Summary statistics of mRNA 3-end sequencing experiments.

| condition | proximal pA site usage | FDR | # | # within LINE repeats proximal | distal |
| --- | --- | --- | --- | --- | --- |
| Matr3 KD1 | up | < 0.05 | 257 | 9 | 6 |
|  |  | >= 0.05 | 417 | 24 | 24 |
|  | down | < 0.05 | 329 | 26 | 5 |
|  |  | >= 0.05 | 428 | 54 | 22 |
|  | neither | - | 2,605 | 150 | 63 |
| Matr3 KD2 | up | < 0.05 | 214 | 11 | 7 |
|  |  | >= 0.05 | 416 | 41 | 27 |
|  | down | < 0.05 | 368 | 14 | 5 |
|  |  | >= 0.05 | 567 | 46 | 26 |
|  | neither | - | 2,256 | 131 | 56 |
| PTB KD | up | < 0.05 | 219 | 5 | 4 |
|  |  | >= 0.05 | 420 | 28 | 21 |
|  | down | < 0.05 | 401 | 18 | 4 |
|  |  | >= 0.05 | 516 | 52 | 28 |
|  | neither | - | 2,332 | 139 | 65 |
| Matr3 KD1 + PTB KD | up | < 0.05 | 370 | 17 | 11 |
|  |  | >= 0.05 | 424 | 34 | 21 |
|  | down | < 0.05 | 399 | 24 | 5 |
|  |  | >= 0.05 | 492 | 44 | 28 |
|  | neither | - | 2,373 | 129 | 59 |

